# Neuro-Cognitive Multilevel Causal Modeling: A Framework that Bridges the Explanatory Gap between Neuronal Activity and Cognition

**DOI:** 10.1101/2023.10.27.564404

**Authors:** Moritz Grosse-Wentrup, Akshey Kumar, Anja Meunier, Manuel Zimmer

## Abstract

Explaining how neuronal activity gives rise to cognition arguably remains the most significant challenge in cognitive neuroscience. We introduce neuro-cognitive multilevel causal modeling (NC-MCM), a framework that bridges the explanatory gap between neuronal activity and cognition by construing cognitive states as (behaviorally and dynamically) causally consistent abstractions of neuronal states. Multilevel causal modeling allows us to interchangeably reason about the neuronal- and cognitive causes of behavior while maintaining a physicalist (in contrast to a strong dualist) position. We introduce an algorithm for learning cognitive-level causal models from neuronal activation patterns and demonstrate its ability to learn cognitive states of the nematode *C. elegans* from calcium imaging data. We show that the cognitive-level model of the NC-MCM framework provides a concise representation of the neuronal manifold of *C. elegans* and its relation to behavior as a graph, which, in contrast to other neuronal manifold learning algorithms, supports causal reasoning. We conclude the article by arguing that the ability of the NC-MCM framework to learn causally interpretable abstractions of neuronal dynamics and their relation to behavior in a purely data-driven fashion is essential for understanding more biological systems whose complexity prohibits the development of hand-crafted computational models.

## 1 Introduction

At least since the work of David Marr (1982), it is widely acknowledged that complex systems can be described at different levels, e.g., at the implementational, the algorithmic, and the computational level. Understanding the neuronal basis of cognition amounts to bridging the explanatory gap between the level of neuronal activity patterns and the level of cognitive states (Changeux and Dehaene, 1989). However, despite the long history of research into this problem, a mathematically rigorous framework for bridging this gap is still lacking. Traditionally, observed statistical associations between single-unit neuronal activities and behaviors have formed the basis for identifying neuronal circuits and hand-crafting mechanistic models of their computations (cf. Borst (2014)). Due to increasing awareness of the importance of (potentially widely distributed) neuronal activity patterns for behavior, and the difficulty in scaling up hand-crafted models to large-scale neuronal recordings, machine learning methods (also referred to as decoding- or multivariate pattern analysis (MVPA) models) have been developed to uncover relations between complex neuronal activity patterns, cognitvie states, and behaviors (Norman et al., 2006; Mitchell et al., 2008; Pereira et al., 2009). More recently, these models have been complemented by algorithms for learning neuronal manifolds that enable the visualization of high-dimensional neuronal dynamics (Mitchell-Heggs et al., 2023) and their relation to behavior (Schneider et al., 2023; Kumar et al., 2023). However, the ability to visualize and decode behavior from complex neuronal activity patterns does not imply that we have revealed their representational contents (Ritchie et al., 2020). Under certain conditions, decoding models can be endowed with a causal interpretation, enabling experimentally testable predictions on causal relations between neuronal activity and behavior (Weichwald et al., 2015). However, we argue that such causal models are not suitable to study relationships between neuronal activity patterns and *cognition* because causal relationships between these two levels would be irreconcilable with physicalism: A causal relation between two variables *X* → *Y* implies that manipulations of *X* can alter the probability of observing specific values of *Y* (Pearl, 2000). This relation is only possible if *X* and *Y* are two separate processes that are linked by a mechanism (cf. Spirtes (2009) for an in-depth discussion of the nature of causal variables). A causal relation between *X* and *Y* thus implies that it must be conceptually possible to intervene on *Y* without intervening on *X*, thereby breaking the mechanism that links the two variables. For instance, if *X* and *Y* represent two different neurons, one could construct an experimental setup that controls the membrane potential of *Y*, thereby eliminating any influence action potentials of *X* can have on the membrane potential of *Y*. If *X* and *Y* were to represent a neuronal- and a cognitive state, however, a causal relation of the form *X* → *Y* would imply that we can (at least conceptually) manipulate the cognitive state without changing the neuronal state that causes it. As such, causal relations between neuronal- and cognitive states would only be meaningful if we adopted a strong dualistic viewpoint in which neuronal- and cognitive states co-existed as independent physical processes. In the present work, we adhere to physicalism (even though our framework may be compatible with a weaker form of *predicate* dualism), i.e., we adopt the view that all mental phenomena are ultimately physical phenomena and hence reject the notion that neuronal activity causes cognition. Instead, we argue in the following that it is reasonable to consider relationships between neuronal activity and cognitive states as *constitutive* or, in more philosophical terms, that cognitive states supervene on neuronal states.

While we consider causal models unsuitable for representing relations between neuronal- and cognitive states, we consider them well-suited to represent relations within each level. We first consider the neuronal level. Building on the example in the previous paragraph, it is perfectly reasonable to consider two neurons as separate entities that can be manipulated individually and that are linked by a physical mechanism. For instance, the sentence *neuron X’s action potential is a cause of neuron Y ‘s membrane potential* is a meaningful causal statement in the sense that it implies an empirically testable prediction that is consistent with our physical understanding of neuronal processes: Manipulating the spiking probability of neuron *X*, e.g., by injecting a current into neuron *X*, will alter the membrane potential of neuron *Y*. We argue that the same holds true for relations between cognitive states. We employ cognitive states to causally reason about our own and other people’s mental processes. For instance, when we say *I am unhappy because I am bored*, we express the causal relation *boredom* → *happiness*. This causal statement implies the empirically testable prediction that if we reduce boredom in a person, we increase their probability of being happy. It is further consistent with the view that *boredom* and *happiness* are two separate processes that can be manipulated independently.

If causal models are well-suited to explain relations within the neuronal and cognitive levels but unsuitable for explaining relations between these two levels, which type of relations should hold between causal models on the neuronal- and causal models on the cognitive level? The solution we advocate here is *causally consistent transformations* between causal models on each level (Rubenstein et al., 2017). Causally consistent transformations are functional mappings between states and interventions of two causal models chosen so that we can reason about the observational effects of causal interventions consistently across both models. For instance, assume the cognitive concepts *boredom* and *happiness* to have the two neuronal realizations ***x*** and ***y*** (with the bold notation indicating that ***x*** and ***y*** may represent elements of high-dimensional sets of neuronal activity patterns), with the causal relations *boredom* → *happiness* and ***x*** → ***y***. Intervening on ***x***, e.g., by electrical stimulation, would then be equivalent to modulating *boredom* and, via the causal effect ***x*** exerts on ***y***, affect *happiness*. In this conceptual framework, which we term *multilevel causal modeling (MCM)*, neuronal cause-effect relations play out in parallel to cognitive cause-effect relations in a consistent manner that enables us to causally reason and explain observations interchangeably on each level. In the MCM framework, neuronal- and cognitive states thus do not co-exist as separate processes but are linked via functional mappings. In other words, neuronal states do not cause but are *constitutive* of cognitive states. The MCM framework thereby provides a theoretical framework to bridge the explanatory gap between neuronal activity and cognition while maintaining a physicalist position.

When attempting to map neuronal- to cognitive states, we must consider that cognitive states are not universal but depend (among other things) on the cultural context (Quinn and Holland, 1987). When attempting to map neuronal- to cognitive states, one question thus seems of particular importance: What is the set of possible cognitive states? Is there, for example, a single cognitive state of being in pain, or should having a headache and feeling back pain be considered two different states? Do all individuals share the same sets of cognitive states? And if so, are the mappings from neuronal to cognitive levels equivalent across subjects? The MCM framework does not assume any fixed set of cognitive states, and we do not argue for the existence of such a universal set. Instead, and in agreement with a recent argument by Krakauer et al. (2017) that neuroscience needs behavior, we let the behavioral context determine the relevant set of cognitive states in a purely data-driven approach.

We illustrate the relationship between cognition, neuronal activity, and behavior in the MCM framework with a (slightly simplified) analogy from physics. The temperature of a gas is proportional to the average kinetic energy of its molecules. To reason about the conditions under which the gas ignites, both levels of description are equivalent, e.g., the two sentences *the gas will ignite if its temperature is raised by T degrees Celsius* and *the gas will ignite if the average kinetic energy of its molecules is increased by K Joules* are causally meaningful and consistent statements. Importantly, many kinetic energy configurations give rise to the same temperature. The gas temperature can not be altered, however, without also changing the kinetic energies. As such, the macroscopic concept of temperature is a causally consistent abstraction of the microscopic kinetic energy configuration with respect to the behavior of the gas. In the MCM framework, the analogies of temperature, kinetic energies, and ignition are cognitive states, neuronal states, and behaviors in the sense that the macroscopic cognitive states are causally meaningful and consistent abstractions of the microscopic neuronal states that cause behavior.

In the following, we introduce the MCM framework in a mathematically rigorous fashion and show how causally consistent transformations between the neuronal- and the cognitive level can be learned from empirical data (Section 2). In Section 3, we illustrate the application of the MCM framework on calcium imaging data from the nematode *C. elegans*. We conclude our work in Section 4 by discussing future extensions and the philosophical implications of the MCM framework.

## 2 Multilevel causal modeling in cognitive neuroscience

We begin this section by discussing the construction of causal models on the neuronal level and elucidating how they are linked to behavior. We then consider the nature of cognitive states and causal models thereof before explaining how causal models on the neuronal- and on the cognitive level can be linked via causally consistent transformations. We conclude the section by discussing how to learn MCMs from empirical data.

### 2.1 Causal models on the neuronal level

To construct causal models on the neuronal level, we first need to decide on a framework for causal modeling. A variety of causal modeling frameworks have been developed for and evaluated on neuronal data (Smith et al., 2011). To be applicable in the MCM framework, we require causal models that can make empirically testable predictions on the effects of experimental interventions. The framework of Causal Bayesian Networks (CBNs) (Pearl, 2000; Spirtes et al., 2000) fulfills this requirement and hence serves as the causal modeling framework in the present setting.

Causal relations in CBNs are modeled by structural causal models (SCMs):

#### Definition 1 (Structural Causal Model – SCM)

*A SCM is a triple* ℳ_***X***_ := {***X, E, F*}** *with* ***X*** *a set of N random variables endogenous to the model*, ***E*** *a set of N exogenous noise variables, and* ***F*** *a set of N functions defining each endogenous variable as a function of its direct causes (i*.*e*., *parents – pa()) and its corresponding exogenous noise variable, so that for each i* ∈ {1, …, *N*} *we have X*_*i*_ := *F*_*i*_(*pa*(*X*_*i*_), *E*_*i*_) *where the F*_*i*_ *are chosen such that no variable is a (direct or indirect) cause of itself*.

The joint probability distribution *P* (***E***) over the exogenous noise variables induces a joint probability distribution *P* (***X***) over the endogenous variables (via the pushforward measure). If the noise variables are mutually independent, the SCM is *causally sufficient*, i.e., no unobserved confounders that influence multiple endogenous variables exist. In the present setting, each endogenous variable *X*_*i*_ represents the activity of one neuron. We note that in general, ***X*** and ***E*** may represent continuous-as well as discrete variables, implying that *P* (***X***) and *P* (***E***) denote probability densities or distributions, respectively. In the following, we assume all variables to be discrete. We consider an extension of the MCM framework to continuous-valued random variables feasible but beyond the scope of the present work. We denote random variables by upper- and their realizations by lower-case letters, e.g., we write *P* (*X*_*i*_ = *x*) to indicate the probability that the neuron represented by the random variable *X*_*i*_ takes on the value *x* ∈ 𝒳.

Causal relations in SCMs are commonly depicted by directed acyclic graphs (DAGs), with an arrow drawn from node *A* into node *B* if the endogenous variable represented by node *A* is a parent of the endogenous variable represented by node *B*. An arrow from *A* into *B* indicates a direct causal influence of *A* on *B*, and a directed path from node *A* to node *B* indicates an indirect causal influence of *A* and *B*. We note that direct- and indirect causal relations are relative to the set of variables endogenous to the model and may change when dropping or adding nodes.

Knowledge of the DAG, in combination with the joint probability distribution over all endogenous variables (represented by the nodes of the DAG), enables us to reason about the probabilistic effects of experimental interventions on variable subsets (Pearl, 2000). Experimental interventions are represented mathematically by the do()-operator, e.g., do(*X*_*i*_ = *x*) represents the experimental intervention of setting variable *X*_*i*_ to the value *x*. In empirical settings, the DAG and the joint probability distribution are usually not known and have to be inferred from a combination of experimental and observational data. We show in Section 2.3 that knowledge of the causal model on the neuronal level is not a prerequisite for constructing a causally consistent mapping between the neuronal- and the cognitive level.

In their original form, SCMs model independent and identically distributed (i.i.d.) data, i.e., data without temporal structure. It is straightforward, however, to extend SCMs to model dynamical neuronal systems by unrolling the endogenous variables across time, as exemplified in the middle row of Figure 1. Here, the set of random variables ***X***[*t*_0_] represents the global state of all *N* neurons at time *t*_0_ and the arrow from ***X***[*t*_0_] into ***X***[*t*_1_] indicates that the global neuronal state at time *t*_0_ is a cause of the global neuronal state at time *t*_1_, in the sense that the state of each neuron at time *t*_1_ is a function of a subset of the neuronal states at time *t*_0_ and their respective exogenous noise term^1^. More formally, we define the extension of SCMs to dynamic settings as follows:

**Figure 1:**
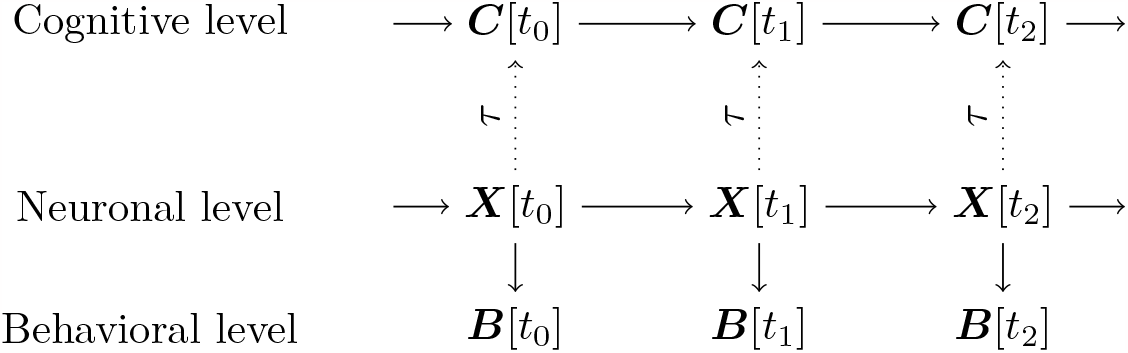
Relations between cognitive-, neuronal-, and behavioral states in MCM. Solid and dotted arrows denote causal and constitutive relations, respectively.

#### Definition 2

*(Dynamic Structural Causal Model – dSCM) A dynamic SCM (dSCM) is family of T triples* ℳ_*X*[*t*]_ := {***X***[*t*], ***E***[*t*], ***F*** [*t*]}, *indexed by t* ∈ {1, …, *T*}, *with* ***X***[*t*] *a set of N random variables endogenous to the model*, ***E***[*t*] *a set of N exogenous noise variables, and* ***F*** [*t*] *a set of N functions defining each endogenous variable as a function of its direct causes (i*.*e*., *parents – pa()) and its corresponding exogenous noise variable, so that for each i* ∈ {1, …, *N*} *and t* ∈ {1, …, *T*} *we have X*_*i*_[*t*] := *F*_*i*_[*t*](*pa*(*X*_*i*_[*t*]), *E*_*i*_[*t*]), *where pa*(*X*_*i*_[*t*]) *may contain any endogenous variables prior to time t*.

We note that we do not allow instantaneous causal relations in Definition 2, i.e., situations where pa(*X*_*i*_[*t*_1_]) may include other neurons at time *t*_1_, because our current understanding of physical processes posits that causes must precede effects. This constraint may have to be reconsidered when modeling dynamical systems that are observed at sampling rates slower than the system’s dynamics. If the exogenous noise terms are mutually independent and pa(***X***[*t*]) only includes variables at the previous time step for all *t*, as in the example in Figure 1, the dSCM represents a first-order Markov process with a transition probability distribution

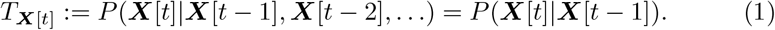

Markov processes are of particular relevance in the MCM framework, because they define causally sufficient state spaces, in the sense that the global state at any given point in time is causally sufficient for the probability distribution over the global state at the next time point. More formally, Markovianity implies (under the faithfulness assumption) that the exogenous noise terms at different time points are mutually statistically independent and hence (via the backdoor criterion, see Pearl (2000)) *P* (***X***[*t*]|***X***[*t* − 1]) = *P* (***X***[*t*]|do(***X***[*t* − 1])) for all *t*, i.e., the observational- and interventional conditional distributions are identical. A counter-example would be the case of an exogenous variable, e.g., a stimulus, affecting both ***X***[*t* − 2] and ***X***[*t*]. In this case, the Markov property would be violated 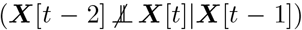 and hence *P* (***X***[*t*]|***X***[*t* − 1]) ≠ *P* (***X***[*t*]|do(***X***[*t* − 1])). As we discuss in Section 2.3, empirical tests for Markovianity are essential to ensure that cognitive variables are causally meaningful. In the following, we assume that the functional mappings between states, as well as the probability distribution over their exogenous variables, do not change over time, i.e., we assume the process to be time-homogeneous. We further assume that any process we consider has converged to a stationary probability distribution. These assumptions are not required from a theoretical perspective but greatly simplify empirical inference.

To model the causal effect of neuronal activity on behavior, we extend the DAG in Figure 1 by a behavioral state vector ***B***[*t*] and let ***X***[*t*] → ***B***[*t*], i.e., we model the behavioral states at time *t* to be caused by (a subset of) the neuronal states at time *t*. We note that it is reasonable to consider relations between neuronal activity and behavior as causal because we can, in principle, manipulate behavior independently of neuronal activity, e.g., we can fixate an animal and thereby prevent neuronal activity from causing an actual movement. We further note that we represent the behavioral states across time by measurement nodes, i.e., by nodes that have no causal effect on any other variables (Markham and Grosse-Wentrup, 2020). As such, the behavioral states do not form a Markov process, and 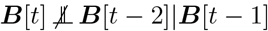 due to the common effect of ***X***[*t* − 2] on both ***B***[*t* − 2] and ***B***[*t*]. We denote the extension of the dSCM by the behavioral state vector as 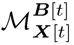.

To model feedback loops of the neuronal system with its environment, the DAG could be further extended by a state vector ***S***[*t*] that represents stimuli, which could be influenced by the system’s past behavior, e.g., ***S***[*t*_0_] → ***X***[*t*_0_] → ***B***[*t*_0_] → ***S***[*t*_1_]. We leave such an extension for future work.

### 2.2 Bridging the neuronal- and the cognitive level

Before discussing how to bridge the neuronal- and the cognitive level, we must first consider the nature of cognitive states. Somewhat surprisingly, there currently exists no generally agreed-upon definition of what a cognitive state is (cf. Prinz (2004), pp. 41 ff.). The broader term *cognition* is commonly used to denote *any kind of mental operation or structure that can be studied in precise terms* (Lakoff and Johnson, 1999). To arrive at a working definition that allows us to operationalize cognitive states for the MCM framework, we consider three refinements of this concept of cognition. First, we propose that a cognitive state must be mathematically quantified to be *studied in precise terms*. Second, we consider an *operation* to refer to a causally meaningful process in the sense that the operation represents a mechanism that translates a system from one state to another. And third, we consider the addendum *or structure* to indicate that a physical substrate supports any mental operation. These considerations lead us to the following working definition, which we will render mathematically precise towards the end of this section:

#### Definition 3 (Cognitive state)

*A cognitive state is a causally meaningful abstraction of a neuronal state*.

To elucidate this definition, we revisit the physical analogy from the introduction. The temperature of a gas is a macroscopic *abstraction* of the microscopic states of the gas molecules because there are infinitely many configurations of kinetic energies of the molecules in the gas that give rise to the same temperature. This relation, however, is asymmetric. It is impossible to change the temperature of a gas without altering the kinetic energies of its molecules. Accordingly, the temperature of a gas and its molecular configuration do not stand in a causal but in a constitutive relationship. The concept of temperature is a *causally meaningful* abstraction because it allows us to reason about the behavior of the gas under experimental interventions, e.g., as in the statement *the gas will ignite if we raise its temperature above a certain threshold*. These interventions are grounded in the microscopic level, i.e., the experimental intervention of raising the temperature is meaningful because there exist microscopic interventions (increasing kinetic energies) that are identical to raising the temperature. Definition 3 expresses the notion that cognitive- and neuronal states stand in a similar constitutive relationship. Consider the cognitive state of being bored. This state is causally meaningful, because, first, being bored can have an effect on other cognitive states, e.g., an experimental intervention that reduces boredom may have the causal effect of increasing happiness, and second, being bored can alter our behavior, e.g., it may lead us to seek out certain (positively stimulating) behaviors. To generalize from this example, the first aspect of Definition 3, that cognitive states are *causally meaningful*, expresses our understanding that we employ cognitive states to reason about mental processes because we consider them to be causally effective. Conversely, we argue that any state that does not stand in a cause-effect relationship to another cognitive state or to behavior is not helpful to reason about mental processes and should therefore be eliminated from a cognitive ontology. To illustrate the second aspect of Definition 3, that cognitive states are *abstractions of neuronal states*, we note that we can represent an organism’s behavior at different levels of granularity. For instance, we may merely distinguish whether an animal is exploring or hunting in its environment, or we may further differentiate the individual movements an animal is performing during the exploration and the hunting processes. In the former case, we would consider all neuronal states that are equally likely to cause an exploratory behavior as constituting one cognitive state. In the latter case, we would enlarge the cognitive state space to be able to distinguish all sets of neuronal states that are equally likely to cause individual movements. In both cases, the cognitive state spaces serve as abstractions of neuronal activity patterns, albeit at different levels of granularity. As in the gas-temperature example, this relationship is asymmetric. There can be many neuronal states that give rise to the same cognitive state. It is impossible to change the cognitive state, however, without also altering the neuronal state. We emphasize that we do not consider the space of cognitive states as constant for a given model system or organism. Instead, we let the granularity at which an organism’s behavior in an environmental context is represented determine the appropriate cognitive state space.

We now formalize the relations discussed above. In analogy to causal models on the neuronal level (cf. Section 2.1), we represent the cognitive state of a model system or organism by a cognitive dSCM ℳ_***𝒞***[*t*]_ := {***C***[*t*], ***E***[*t*], ***F*** [*t*]}, with ***C***[*t*] the cognitive state vector (note that ***E***[*t*] and ***F*** [*t*] are specific to each model and not shared across neuronal- and cognitive dSCMs). To link the the neuronal- and the cognitive dSCM in a causally consistent manner, we require the two models to be *behaviorally* and *dynamically* causally consistent:

#### Definition 4 *(Behavioral Causal Consistency – BCC)*

*Let* 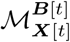 *a neuronal dSCM in a behavioral context and* ℳ_***𝒞***[*t*]_ *a cognitive dSCM. Denote the state space of* ***X***[*t*] *and* ***C***[*t*] *by* ***𝒳*** *and* ***𝒞***, *respectively. We call the triple* {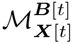, ℳ_***𝒞***[*t*]_, *τ* } *behaviorally causally consistent if τ* : ***𝒳*** ↦ ***𝒞*** *is a surjective mapping such that for all* ***b*** ∈ ℬ *and for all* ***x*** ∈ ***𝒳***, ***c*** ∈ ***𝒞*** *with* ***c*** = *τ* (***x***) *we have*

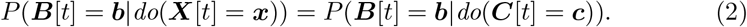

If a neuronal- and a cognitive SCM are behaviorally causally consistent, every experimental intervention on the neuronal level has a matching intervention on the cognitive level, in the sense that both interventions lead to the same probability distribution over the behaviors. The surjectivity of *τ* ensures, first, that the cognitive level is an abstraction of the neuronal level, i.e., many neuronal states may map to the same cognitive state, yet distinct cognitive states can only be linked to distinct neuronal states, and, second, that every cognitive state has at least one neuronal representation. Intuitively, the cognitive SCM constitutes a lossless compression of all information in the neuronal SCM that is causally relevant for a given set of behaviors. As such, behavioral causal consistency allows us to reason interchangeably about the causes of behaviors on the neuronal- and on the cognitive level. Importantly, behavioral causal consistency endows cognitive states with a causal meaning, in the sense that cognitive interventions, for which it may be unclear how the intervention can be experimentally implemented (e.g., *increase happiness*), can be translated into equivalent interventions on the neuronal level, for which a well-defined experimental procedure is available (e.g., *stimulate a set of neurons*).

#### Definition 5 *(Dynamic Causal Consistency – DCC)*

*We call a triple* {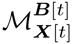, ℳ_***C***[*t*]_, *τ* } *dynamically causally consistent if for all pairs of cognitive states* ***c, c***′ ∈ ***𝒞***

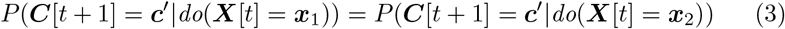

*for all* ***x***_1_, ***x***_2_ ∈ *τ* ^*−*1^(***c***) *where τ* ^*−*1^(***c***) *is the pre-image of* ***c***.

*In this case, we define do*(***C***[*t*] = ***c***) := *do*(***X***[*t*] = ***x***|***x*** ∈ *τ* ^*−*1^(***c***)) *and have*

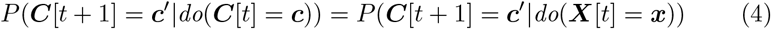

*for all* ***c, c***′ ∈ ***𝒞*** *and* ***x*** ∈ *τ* ^*−*1^(***c***).

While behavioral causal consistency defines the consistency of neuronal- and cognitive states with respect to behavior, dynamic causal consistency defines consistency with respect to the dynamics of the neuronal system. Intuitively, a cognitive SCM that is DCC constitutes a lossless compression of all information in the neuronal SCM that is causally relevant to the dynamics of the neuronal system. In analogy to behavioral causal consistency, DCC grounds causal relations between cognitive states in the neuronal level, e.g., statements such as *reducing boredom is likely to lead to increased happiness* can be translated into the neuronal-level statement *inducing a neuronal activity pattern* ***x***′ *that represents low boredom, e*.*g*., *by electrical stimulation, is likely to lead to a neuronal state that corresponds to increased happiness*.

We are now in a position to formally introduce the primary contribution of this work:

#### Definition 6 *(Neuro-Cognitive Multilevel Causal Model – NC-MCM)*

*We call a triple* {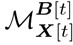, ℳ_***C***[*t*]_, *τ* } *a neuro-cognitive multilevel causal model (NC-MCM) if it is behaviorally and dynamically causally consistent*.

We remark that we have chosen *τ* not to depend on time because we consider a constant mapping between levels more desirable. It would be straightforward to extend the definition to time-varying mappings, but this generalization would substantially complicate learning the mapping from empirical data. We further note that in contrast to Rubenstein et al. (2017) we only consider experimental interventions on the full neuronal- and cognitive state vectors at each time step and leave the extension of NC-MCMs to interventions on subsets of state variables for future work.

We illustrate the concept of a NC-MCM in the following toy example.

**Example 1** *Consider a neuronal dSCM* 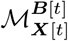 *with one endogenous variable (N* = |***X***| = 1*) and discrete state space* 𝒳 = {*x*_1_, *x*_2_, *x*_3_, *x*_4_}, *where each state represents one out of a total of four neuronal activity patterns. Further, assume there are three distinct behaviors* ℬ = {*b*_1_ (*e*.*g*., *move*), *b*_2_ (*e*.*g*., *feed*), *b*_3_ (*e*.*g*., *rest*)} *with conditional probabilities given the neuronal state*

| $P(\mathbf{B}\vert \mathbf{X})$ | $x_1$ | $x_2$ | $x_3$ | $x_4$ |
| --- | --- | --- | --- | --- |
| $b_1$ | 0.5 | 0.5 | 0.1 | 0.1 |
| $b_2$ | 0.4 | 0.4 | 0.1 | 0.1 |
| $b_3$ | 0.1 | 0.1 | 0.8 | 0.8 |

*Note that the conditional probability distribution over the three behaviors is identical for* {*x*_1_, *x*_2_} *and for* {*x*_3_, *x*_4_}. *As such, one may interpret the system as having two distinct macro-states*, {*x*_1_, *x*_2_} *and* {*x*_3_, *x*_4_}, *with each macro-state subserving a particular behavioral pattern that is expressed by a constant probability distribution over the three behaviors. We may thus define a mapping*

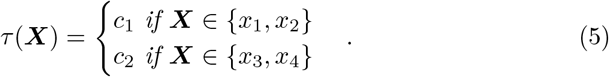

*These two macro-states form the cognitive state space 𝒞* = *c*_1_, *c*_2_, *where we may choose to name the two cognitive states foraging and recuperating, respectively. Next, assume the neuronal dSCM has the steady-state transition probability distribution T*_***X***_ *shown in the left-hand side of Figure 2. The transition matrix T*_***X***_ *and the mapping τ induce a cognitive dSCM with the transition probabilities shown in the right-hand side of Figure 2. The triple* {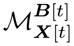, ℳ_***C***[*t*]_, *τ*} *is behaviorally causally consistent, because the cognitive macro-states condense all information in the neuronal micro-states that are causally relevant for the set of behaviors, i*.*e*., *for every intervention on the neuronal micro-level there is an equivalent intervention on the cognitive macro-level that entails the same probabilities over the behaviors (cf. Definition 4). The triple is further dynamically causally consistent because the probabilities of remaining in the current or progressing to another cognitive state are identical for each neuronal state that maps to the same cognitive state. Because the triple is behaviorally and dynamically causally consistent, it fulfills the criteria for a NC-MCM. As such, the cognitive dSCM is a causally meaningful abstraction of the underlying neuronal dSCM because causal statements on the effects of current cognitive states on behavior and on future cognitive states are grounded in the neuronal level, e*.*g*., *the cognitive-level statement ‘the animal is moving around and feeding because it is foraging’ can be translated into the neuronal-level statement ‘the animal is moving around and feeding because it is in neuronal state x*_1_ *or x*_2_*’. We note that behavioral- does not imply dynamic causal consistency, e*.*g*., *introducing an asymmetry in the neuronal-level transition probabilities in Figures 2 would break dynamic- but not behavioral causal consistency*.

**Figure 2:**
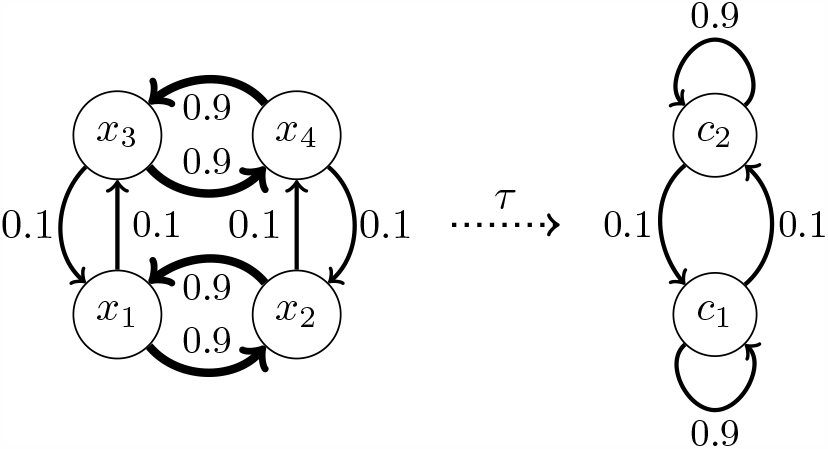
State-space diagram with transition probabilities for Example 1.

We remark that the toy example above is constructed to illustrate the idea that in the NC-MCM framework redundancies in the neuronal dynamics and their relation to behaviors are exploited to build a cognitive-level model that preserves all causally relevant information. In practice, the challenge is to learn which (potentially widely-distributed and complex) neuronal activity patterns are redundant with respect to a behavioral context *and* with respect to the neuronal dynamics, a problem that we turn to in the next section.

### 2.3 Learning neuro-cognitive multilevel causal models

A NC-MCM model is specified by the triple {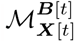, ℳ _***C***[*t*]_, *τ* }, a neuronal-level dSCM in a behavioral context, a cognitive-level dSCM, and a causally consistent mapping between the two. A learning problem arises when any subset of these three components is not fully specified and needs to be inferred from data. In this section, we consider the case where we have access to a set of samples 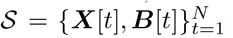, generated by some unknown neuronal dSCM in a behavioral context, and our goal is to learn a mapping *τ* that induces a (behaviorally and dynamically) causally consistent cognitive dSCM. We consider this setting of particular interest because it amounts to learning the cognitive states that a neuronal system has developed in a specific behavioral context from empirical data. Other learning problems are briefly discussed in Section 4.

Learning a mapping *τ* that gives rise to a causally consistent cognitive dSCM proceeds in two steps. The first step is to learn a causal model between neuronal states and behaviors. This model then forms the basis for learning the mapping *τ* that induces a behaviorally and dynamically consistent cognitive dSCM. In this section, we discuss these steps from a conceptual perspective. A particular instantiation of a pipeline that learns a causally consistent cognitive dSCM from empirical data S is presented in Section 3.2.

Learning a causal model that relates neuronal activity to behavior amounts to learning the interventional distribution *P* (***B***[*t*]|do(***X***[*t*])). The gold standard to identify interventional distributions is by experimentation, i.e., by repeatedly setting ***X***[*t*] to random values via an intervention and observing the induced behaviors. However, large-scale neuronal stimulation with concurrent recordings of neuronal activity and behavior remains a challenge (Yazdan-Shahmorad et al., 2016). Alternatively, we may attempt to learn the interventional distribution from observational data only. A variety of causal inference algorithms have been developed for this purpose (Pearl, 2000; Spirtes et al., 2000; Peters et al., 2017) and applied to neuronal data (Smith et al., 2011; Weichwald et al., 2015; Grosse-Wentrup et al., 2016). However, causal structure learning from observational data is also a hard problem, the computational complexity of which typically grows exponentially in the number of variables. We thus follow a third approach in which we merely learn an observational prediction model *P* (***B***[*t*]|***X***[*t*]) from the set of samples 𝒮. We discuss the implications of substituting an interventional model for an observational one at the end of this section.

After learning the interventional or observational distribution between neuronal activity and behavior, the second step is to learn the actual mapping *τ*. Learning this mapping can be broken down into two further steps. The first step is to construct a behaviorally consistent mapping between neuronal activity and cognitive states. The second step is to empirically test whether the induced cognitive dSCM is also dynamically causally consistent. We note that designing a one-step algorithm that directly constructs a mapping that is guaranteed to be behaviorally and dynamically consistent would be preferable but is beyond the scope of the present work.

To construct a behaviorally consistent mapping, we need to find subsets of the neuronal feature space, i.e., partitions, for which *P* (***B***[*t*]|***X***[*t*]) is constant. One way to solve this problem is to first learn a model that predicts the probability of each behavior as a function of the neuronal state, e.g., using a logistic regression model, a neural network model, or any other suitable modeling approach. In the second step, the predicted probabilities can then be clustered, e.g., using k-means or any other preferred clustering algorithm. The inferred clusters define a partition of the neuronal feature space with (approximately) constant conditional probabilities over all behaviors, i.e., the mapping *τ*. This mapping then induces a cognitive dSCM that is by construction behaviorally consistentm, with the number of cognitive states being equal to the number of clusters.

Because behavioral does not imply dynamic causal consistency, we additionally need to test whether the learned mapping *τ* also induces a dynamically causally consistent cognitive dSCM. To do so, we first note that our definition of dynamic causal consistency (Def. 5) is identical to condition (3) in Theorem 1 of Burke and Rosenblatt (1958), with the exception that the former and the latter are based on interventional- and observational transition probabilities, respectively. Accordingly, we term condition (3) in Burke and Rosenblatt (1958) dynamic *observational* consistency. We can then invoke Theorem 4 of Burke and Rosenblatt (1958) to conclude that Markovianity of the cognitive dSCM is sufficient for dynamic observational consistency. Finally, we recall from Section 2.1 that Markovianity of a dSCM implies that the observational- and interventional transition probabilities coincide. As such, Markovianity of the cognitive dSCM is also sufficient for dynamic causal consistency.

To empirically test a cognitive dSCM for Markovianity, we test the null-hypothesis *H*0 : ***C***[*t* − 1] ⊥ ***C***[*t* + 1] |***C***[*t*]. If we find sufficient evidence against Markovianity, we reject the null hypothesis and conclude that the cognitive dSCM is not dynamically causally consistent. Otherwise, we accept the null hypothesis and conclude that the cognitive dSCM induced by *τ* is behaviorally and dynamically causally consistent with the data-generating neuronal dSCM. We note that if we find a cognitive dSCM not to be dynamically causally consistent, we can vary the number of clusters to tune the granularity of the cognitive dSCM and repeat the test for Markovianity.

If we base the procedure described above on the observational distribution *P* (***B***[*t*]|***X***[*t*]) (instead of on the interventional distribution *P* (***B***[*t*]|do{***X***[*t*]})), the cognitive dSCM is behaviorally *observationally* consistent (BOC) but not (in general) behaviorally causally consistent (BCC). However, the *causal coarsening theorem* states that a causal partitioning is almost always a coarsening of an observational partitioning (Chalupka et al., 2015, 2017). As such, a cognitive dSCM that is BCC can be obtained from a cognitive dSCM that is BOC by fusing subsets of cognitive states. While experimental interventions may be required to identify which cognitive states cause behaviors with identical probabilities and thus should be fused, the number of required interventions is on the order of the number of cognitive states and not on the order of the (potentially orders of magnitude larger) number of neuronal states. As such, learning a cognitive dSCM on observational data first and reducing the number of cognitive states by experimental interventions afterward is experimentally more tractable than directly constructing the interventional distribution *P* (***B***[*t*]|do(***X***[*t*])) on the neuronal level. If even a reduced number of experimental interventions is not feasible, we may have to accept that a cognitive dSCM is merely BOC.

## 3 Multilevel causal modeling in *C. elegans*

In this section, we demonstrate the application and utility of the NC-MCM framework on calcium imaging data recorded in the nematode *C. elegans*. After introducing the data in Section 3.1, we show how to learn cognitive dSCMs that are behaviorally observationally consistent and test them for dynamic causal consistency (Section 3.2). In Section 3.3, we study the NC-MCMs in terms of the behavioral motifs they uncover (Section 3.3.1), the representations of *C. elegans*’ neuronal manifold as a graph (Section 3.3.2), and the insights these models enable into decision making in *C. elegans* (Section 3.3.3).

*C. elegans* is an ideal model system to demonstrate the utility of the NCMCM framework for multiple reasons. It has only 302 neurons, about a third of which can be simultaneously recorded by Ca^2+^ imaging in an individual animal at single-cell resolution (Schrödel et al., 2013; Venkatachalam et al., 2016; Nguyen et al., 2016). Due to the stereotypical nature of the nematode’s neuronal system, many of these neurons can be identified and thus compared across animals. *C. elegans* further has a small behavioral repertoire of motor programs like, e.g., forward crawling, backward crawling and turning. This moderate complexity, in combination with extensive prior knowledge of relations between neuronal activity and behavior, allows us to demonstrate the neuropysiological plausibility of the insights derived from the NC-MCM framework, paving the way for its application to more complex model systems.

We remark that the framework for learning a causally consistent cognitive dSCM is model agnostic in the sense that each of the computational steps in Section 2.3 can be accomplished by different algorithms that solve the same computational task, e.g., the behavioral probabilities can be estimated from the neuronal states via logistic regression, (deep) neuronal networks, or any other suitable decoding method. We follow the general guideline that simple models should be preferred over complex ones if they provide sufficiently accurate results. The pipeline described below is implemented as the function *learn mcm(neuronal data, behavioral labels)* in the *nilab* toolbox available for the *Julia* programming language at https://github.com/moritzgw/nc-mcm. The code to reproduce the empirical results presented here, which is based on the *nilab* toolbox and customized *Python* functions for plotting, is also available in that repository. We note that due to the inherent randomness of some of the modules, e.g., *k*-means clustering and permutation-based statistical tests, results may slightly vary across multiple runs. We carefully checked that the empirical results reported below are qualitatively consistent across runs.

### 3.1 The data

The data set we use has been recorded by Kato et al. (2015) and is available at https://osf.io/2395t/. We use the data subset that has been recorded without externally applied stimulation. It consists of data from five immobilized worms with 107 – 131 neurons recorded in each individual worm for a period of 18 minutes at a sampling rate of approximately 2.85 Hz. We subsequently refer to a sample of the calcium traces as the neuronal state vector ***X***[*t*], which represents the state of all neurons recorded in one animal at time *t*.

Even though all five worms were immobilized during the recordings, established equivalences between the activity of individual neurons and the worms’ behavior were used to label each neuronal state vector with the behavior that would have been concurrently observed in a non-immobilized worm; these encompass motor commands for forward crawling, forward slowing, backward crawling, turning dorsally and turning ventrally (Kato et al., 2015). Based on distinct activity patterns, the neuronal state corresponding to backward crawling was further subdivided into reversal 1, reversal 2, and sustained reversal (Kato et al., 2015). To date, the behavioral correlates of these sub-divisions are not known. The behavioral labels assigned to the neuronal states by Kato et al. (2015) are ℬ = {reversal1 (*rev1*), reversal2 (*rev2*), slowing (*slow*), ventral turn (*vt*), sustained reversal (*revsus*), forward movement (*fwd*), dorsal *turn* (*dt*)}. Neuronal states that appeared ambiguous, i.e., for which the behavioral label could not be clearly identified, are labeled as *nostate*, corresponding to a small fraction (∼ 1%) of the data frames.

### 3.2 Learning a cognitive model of *C. elegans*

Learning a cognitive model proceeds in three steps: Predicting the probabilities of all behaviors for every neuronal state vector, clustering the predicted probabilities to obtain the cognitive states, and testing the induced cognitive state transitions for Markovianity.

To learn an observational model that relates neuronal activity patterns to behavior in each individual animal, i.e., to estimate *P* (***B***|***X***), we trained a set of logistic regression models to discriminate between all pairs of the eight behaviors, resulting in 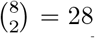 models per animal. We chose logistic regression models for this step because they directly estimate the probability of a behavior as a function of the neuronal state. Each classifier was trained on all recorded neurons of each individual animal, excluding only those neurons that were used to identify the behavioral labels (AVAL, AVAR, SMDDR, SMDDL, SMDVR, SMDVL, RIBR, and RIBL) (Kato et al., 2015). Classification accuracy was evaluated via majority voting in combination with 10-fold cross-validation using random data splits. This evaluation resulted in a group-average decoding accuracy of 92.2%, with decoding results between 89.4% and 95.6% across individual worms, indicating that the linear logistic regression models provide a good characterization of the observational relationships between neuronal activities and behaviors. For subsequent modeling steps, the regression models were trained on all data without cross-validation. We then used these models to compute the conditional probabilities each pairwise classifier assigned to their respective behaviors as a function of the neuronal state, resulting in a 28-dimensional vector of behavioral probabilities for each neuronal state, which we subsequently term the behavioral probability trajectories. We note that the logistic regression models estimate observational- and not interventional probabilities (cf. Section 2.3).

To learn the mapping *τ*, we employed *k*-means clustering in the 28-dimensional space of probability estimates to assign neuronal states to clusters with (approximately) constant conditional behavioral probabilities (cf. Definition 4). We varied the number of clusters for *k*-means between two and 20 and re-ran *k*-means 100 times for each *k* with random initial seeds. For each *k* and run, we thereby obtained an assignment of every neuronal state to one out of *k* cognitive states, which we subsequently refer to as a cognitive state trajectory ***C***[*t*].

We then tested each of the 5 (worms) × 20 (range of the number of cognitive states) × 100 (clustering runs) cognitive state trajectories for Markovianity by, first, estimating the first- and second-order cognitive state transition probability matrices *P* (***C***[*t*]|***C***[*t* − 1]) and *P* (***C***[*t*]|***C***[*t* − 1], ***C***[*t* − 2]), second, computing the total variance of *P* (***C***[*t*]|***C***[*t* − 1]) across all states of ***C***[*t* − 2] (whose expected value for a Markov process is zero), and third, computing the same total variance for one thousand simulated Markovian cognitive state trajectories with state transition probability matrix *P* (***C***[*t*]| ***C***[*t* − 1]). This enabled us to estimate the *p*-value under the null hypothesis of a Markovian cognitive state trajectory as the frequency at which the simulated total variance exceeded the observed one^2^. Because higher *p*-values signify a better clustering result, in the sense that the learned cognitive state trajectory exhibits no evidence against Markovianity (cf. Section 2.3), we then picked, for each worm and number of cognitive states, the cognitive state trajectory with the highest *p*-value^3^. Figure 3 shows the *p*-values of the selected cognitive state trajectories as a function of the number of cognitive states for each worm (dots) as well as averaged across all worms (line). When choosing *α* = 0.05 as a *lower* threshold for *accepting* the null hypothesis, we obtain Markovian cognitive state trajectories for any number of cognitive states in the range from three to 19, with a maximum average *p*-value for seven states. As such, we infer that all cognitive state trajectories with three to 19 states are dynamically causally consistent. As discussed below, we can then control the degree of granularity at which we study the dynamics of *C. elegans* by varying the number of cognitive states in this range.

**Figure 3:**
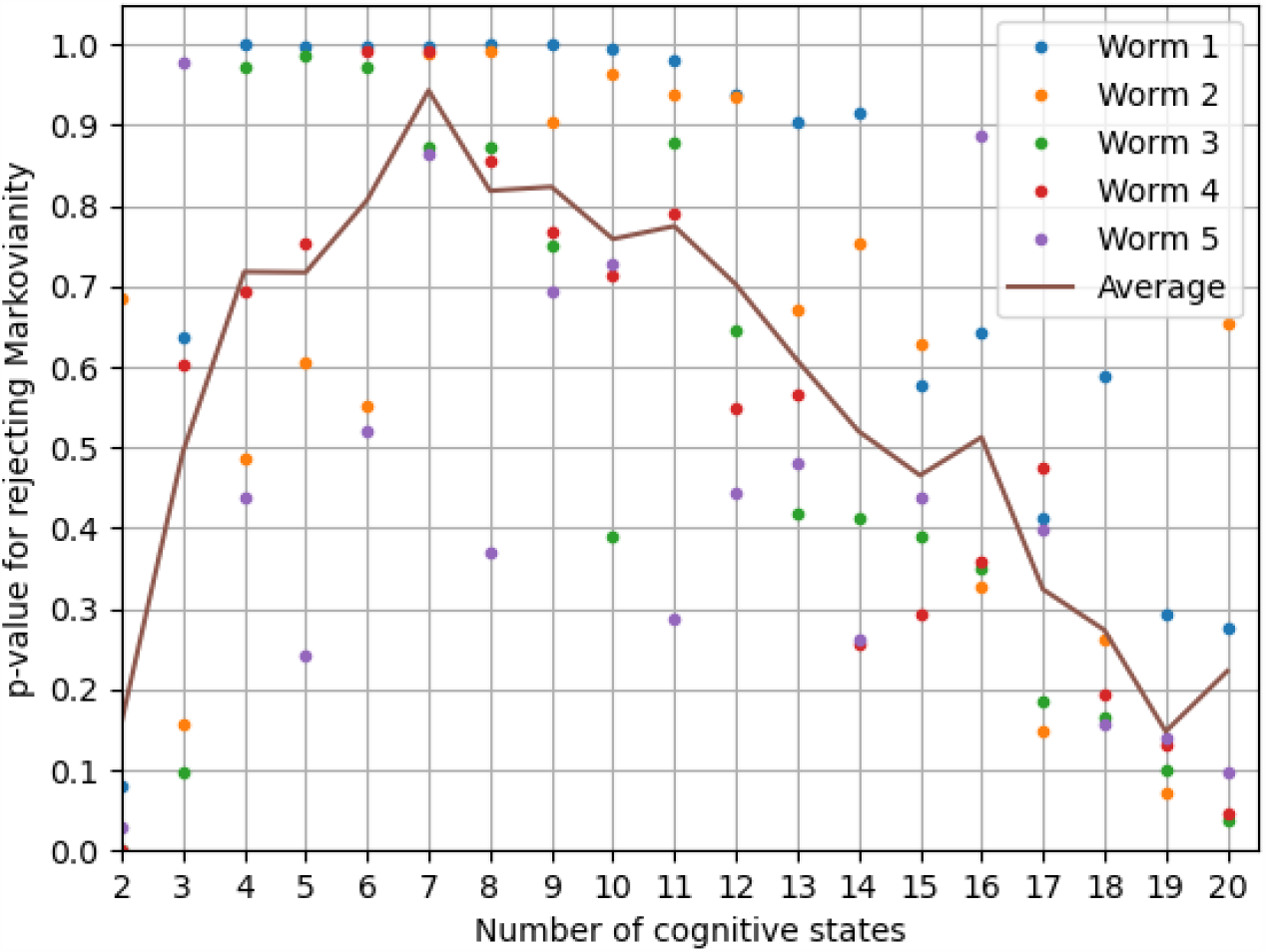
*p*-values for rejecting the null-hypothesis of Markovianity for each worm (and averaged across worms) as a function of the number of cognitive states.

To build a better intuition for the procedure of learning a cognitive state trajectory, Figure 4 illustrates the relevant steps on the data of the first worm with five cognitive states (all trajectories are projected to their first two principal components for visualization): The neuronal state trajectories together with the behavioral labels (A) are used to estimate the behavioral probability trajectories (B). These trajectories are then clustered to assign each element of the behavioral probability trajectories to a cognitive state (C). Re-projecting the cognitive labels onto the neuronal state trajectories gives the neuro-cognitive state trajectories (D), from which we can compute the cognitive state transition model *P* (***C***[*t*]|***C***[*t* −1]).

**Figure 4:**
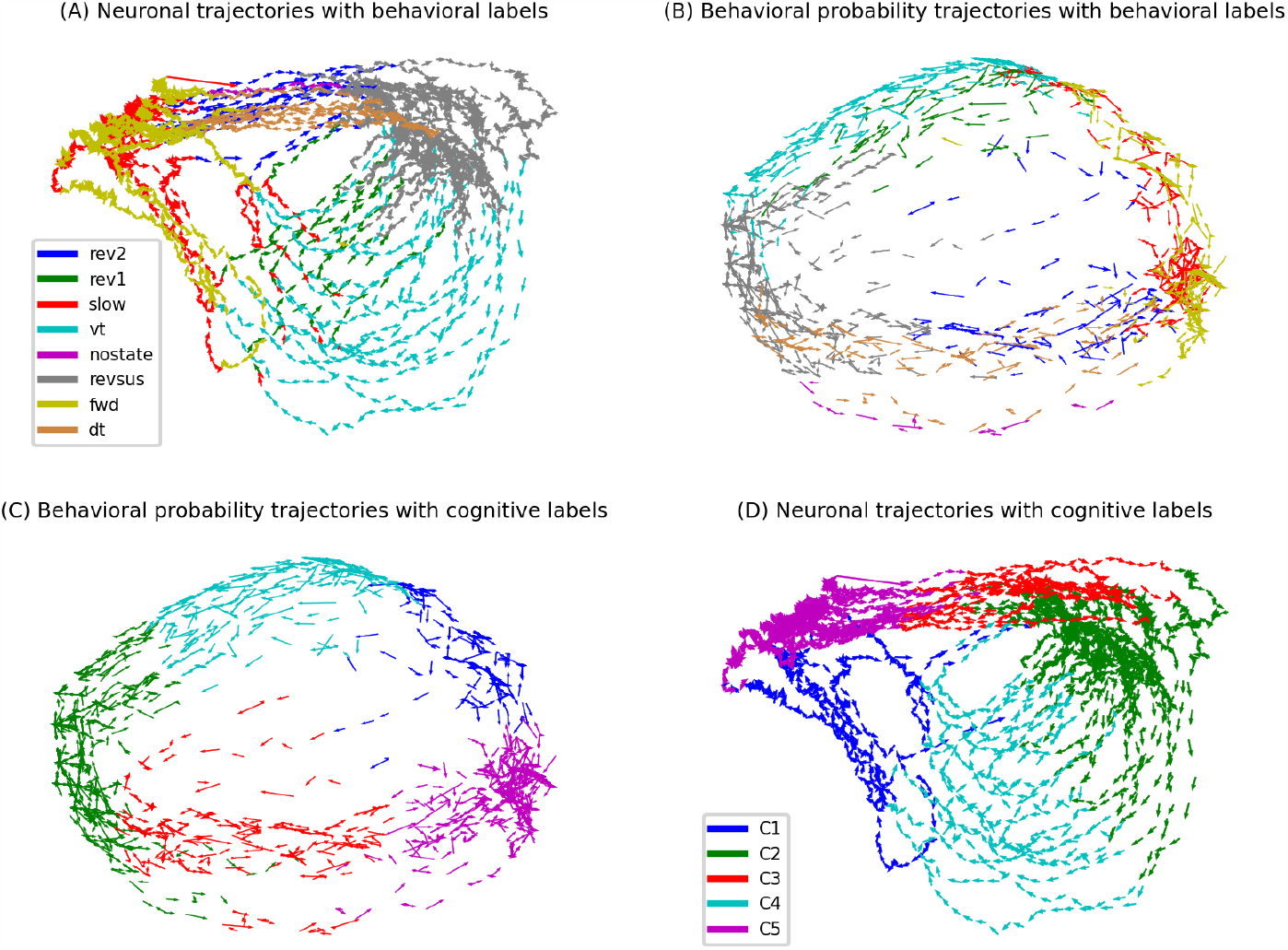
Illustration of the procedure of learning a cognitive state trajectory (see main text for details).

We remark that the cognitive dSCM of each animal is behaviorally observationally consistent by construction and dynamically causally consistent if the cognitive dSCM is Markovian^4^. Because we built the cognitive dSCM on the observational distribution *P* (***B***|***X***) rather than on the interventional distribution *P* (***B***|do(***X***)), which is not available in the present setting, behavioral causal consistency is not guaranteed (cf. Section 2.3). In its present form, the cognitive dSCM thus only supports causal statements about the dynamics of the cognitive dSCM but not about its relation to behavior. However, its observational behavioral consistency can be used to derive causal hypotheses that guide the design of experimental studies to establish causal relations between neuronal states and behaviors via interventions, a topic we discuss in Section 3.3.3.

### 3.3 Interpreting the cognitive model of *C. elegans*

In this section, we demonstrate the ability of the learned cognitive models to reveal behavioral motifs of *C. elegans* (Section 3.3.1), show that from a data-visualization perspective, the NC-MCM framework can be understood as a neuronal manifold learning technique that abstracts the essential features of a manifold into a directed graph (Section 3.3.2), and discuss how the cognitive model can provide insights into decision-making processes (Section 3.3.3).

#### 3.3.1 Behavioral motifs of *C. elegans*

In the following, we juxtapose the behavioral state transition diagram with the state diagram that is obtained by expanding the behavioral- by the cognitive states, i.e., we compare *P* (***B***[*t*)|***B***[*t* − 1]) and *P* (***C***[*t*], ***B***[*t*)|***C***[*t* − 1], ***B***[*t* − 1]). In particular, we show that the behavioral state transition diagram conflates distinct behavioral motifs that are revealed when considering the behavior commands in relation to the cognitive states in which they appear. We first illustrate the results on the third worm in the data set because this worm exhibits the most simple state transitions of all the five worms. We then show that the behavioral motifs discussed below are present with small variations in every worm.

Figure 5 shows the behavioral- (A, left column) and the cognitive-behavioral state diagram with three cognitive states (B, right column) of the third worm in the data set^5^. In the left diagram, each node represents one behavioral state. The size of each node represents the probability of the worm being in the corresponding behavioral state at any point in time. The width of an outgoing edge represents the probability of the worm transitioning from a particular state to another state (edges indicating the probabilities of staying in a particular behavioral state, i.e., edges from one node to itself, are not drawn to reduce clutter)^6^.

**Figure 5:**
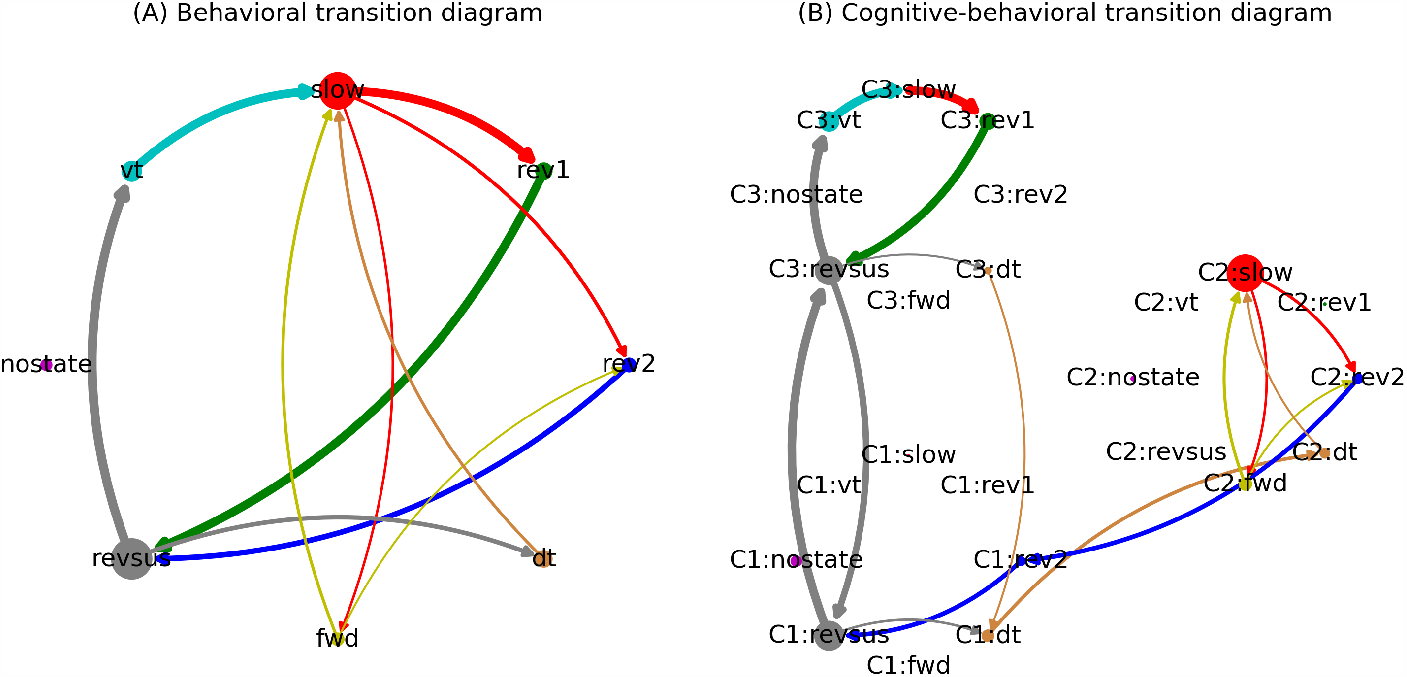
Behavioral- and cognitive-behavioral state diagrams of the third worm (see text for details). Arrows that account for less than 0.1% of outgoing transitions of each node have been removed to reduce clutter.

Figure 5.A reveals a clear structure in the worm’s behavioral command (or states) sequences. The worm is most often found executing a sustained reversal state (*revsus*). A sustained reversal is either followed by a ventral- (*vt*) or dorsal (*dt*) turn state, both of which segue into a slowing state (*slow*). The slowing state alternates with a forward state (*fwd*) before returning to a sustained reversal state via either of two types of reversal states (*rev1* or *rev2*).

We have chosen observational language to describe this behavioral state transition diagram because the behavioral dynamics are not Markovian (the statistical test for Markovianity described in Section 3.2 rejects the null hypothesis of Markovianity at *α* = 0.05 (*p* = 0.015)). This non-Markovianity indicates that past behavioral states provide information about the future state that is not contained in the current state, as was suggested by Kato et al. (2015). This fact becomes intuitively plausible when considering the cognitive-behavioral state transition diagram in Figure 5.B. This diagram expands the behavioral by the cognitive states, i.e., we represent the eight behaviors separately for each cognitive state in which they occur^7^. This Markovian representation reveals two distinct behavioral motifs: A *revsus* → *vt* → *slow* → *rev1* → *revsus* and a *revsus* → *dt* → *fwd* ↔ *slow* → *rev2* → *revsus* loop. These two loops that occur in cognitive states C3 and C2, respectively, are connected via cognitive state transitions between C1 and C3 during a sustained reversal (which we will discuss in more detail in Section 3.3.3). The first motif represents a repeated execution of a state sequence composed of sustained reversal followed by a ventral turn, slowing and reversal 1. The second motif is composed mostly of switching between slowing and forward crawling states. In the third worm, this motif always starts with a dorsal turn state and terminates with a reversal 2. These behavioral motifs are well known in *C. elegans* (Pierce-Shimomura et al., 1999; Kimura et al., 2010). The first motif corresponds to a pirouette, where animals execute multiple reverse-turning events in close succession. The second motif corresponds to a forward run episode, where animals persistently head in a forward direction. This motif is terminated by a rev2 state, corresponding to the neuronal signature that terminates a forward run episode (cf. Kato et al. (2015)).

Notably, the NC-MCM framework discovered these motifs without any prior knowledge of the underlying behavioral dynamics. Rather, the behavioral motifs are revealed by the NC-MCM framework because they are embedded in distinct neuronal dynamics. Importantly, this entails the ability of the NC-MCM framework to assign time frames that were manually annotated with the same behavioral label to distinct cognitive states if the executed behaviors differ in their neuronal realization. In Figure 5, this is apparent in two distinct slowing behaviors, one in each behavioral motif, that are supported by cognitive states C2 and C3, respectively. In C2, slowing could correspond to brief episodes of reducing locomotion speed while the animals remain in a forward run (Kato et al., 2015). In C3, slowing could correspond to intermittent pausing or brief forward crawling during successive re-orientations. In the behavioral state diagram in the left column of Figure 5, these two different types of slowing movements are conflated. This conflation leads to a non-Markovian representation because the probabilities of entering a *rev1* and *rev2* state are higher when the slowing movement is preceded by a ventral and a dorsal turn, respectively. Conversely, the Markovian representation of the cognitive-behavioral state diagram ensures that it correctly distinguishes seemingly similar behaviors that are supported by distinct neuronal dynamics.

While the third worm exhibits the most distinct separation of the two behavioral motifs, both motifs are present in every worm (Figure 6). Because the worms slightly vary in the complexity of their state transitions, with the other worms also showing infrequent state transitions between the two motifs, the number of cognitive states at which the two behavioral motifs first emerge varies across worms (four cognitive states for the first two worms and three cognitive states for the remaining three worms).

**Figure 6:**
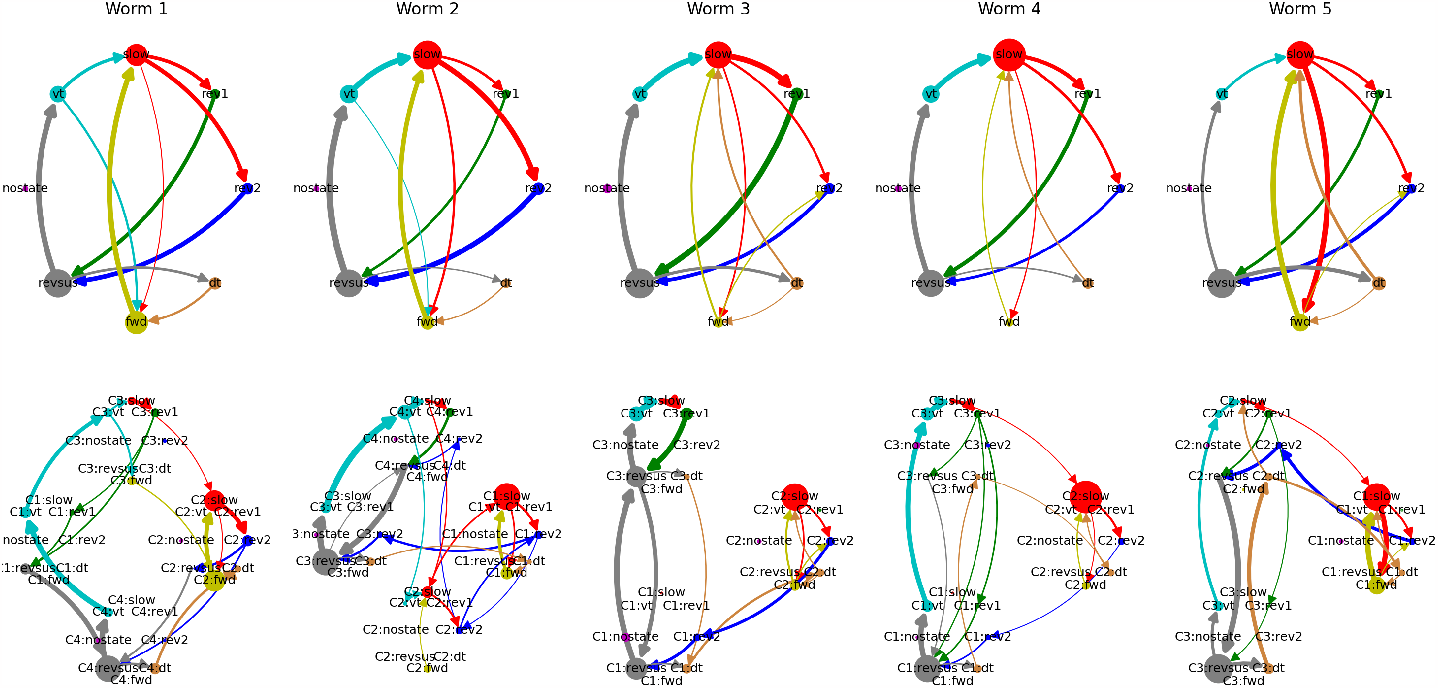
Behavioral- and cognitive-behavioral state diagrams of all worms. Arrows that account for less than 0.075% of outgoing transitions of each node have been removed to reduce clutter.

#### 3.3.2 Representing *C. elegans*’ neuronal manifold as a directed graph

Individual neurons are embedded in brain networks that collectively organize their high-dimensional neuronal activity patterns into lower-dimensional neuronal manifolds (Churchland et al., 2012; Gallego et al., 2017). The goal of neuronal manifold learning techniques is to find low-dimensional representations of neuronal data that enable insights into the structure of neuronal dynamics and their relation to behavior (Mitchell-Heggs et al., 2023). In neuroscience, classic dimensionality reduction techniques, such as principal component analysis (PCA), Laplacian eigenmaps (LEM), and t-SNE, are complemented by modern deep learning techniques such as CEBRA (Schneider et al., 2023) and BundDLe-Net (Kumar et al., 2023). In the following, we show that the cognitive-behavioral state diagrams introduced in the previous section can be interpreted as a neuronal manifold learning technique that represents the essential aspects of the manifold as a directed graph.

Figure 7 shows a side-by-side comparison of the neuronal manifold of the third worm as inferred by BundDLe-Net (Kumar et al., 2023) (left column) and the cognitive-behavioral state diagram of the NC-MCM framework (right column)8. Looking at the left column, it is immediately apparent that the neuronal manifold exhibits two main cycles, a *revsus* (gray) → *vt* (turquoise) → *slow* (red) → *rev1* (green) → *revsus* (gray) and a *revsus* (gray) → *dt* (orange) → *slow* (red) → *fwd* (yellow) → *rev2* (blue) → *revsus* (gray) cycle, that correspond to the behavioral motifs revealed by the cognitive-behavioral state diagram in the right column of Figure 7 and discussed in the previous section.

**Figure 7:**
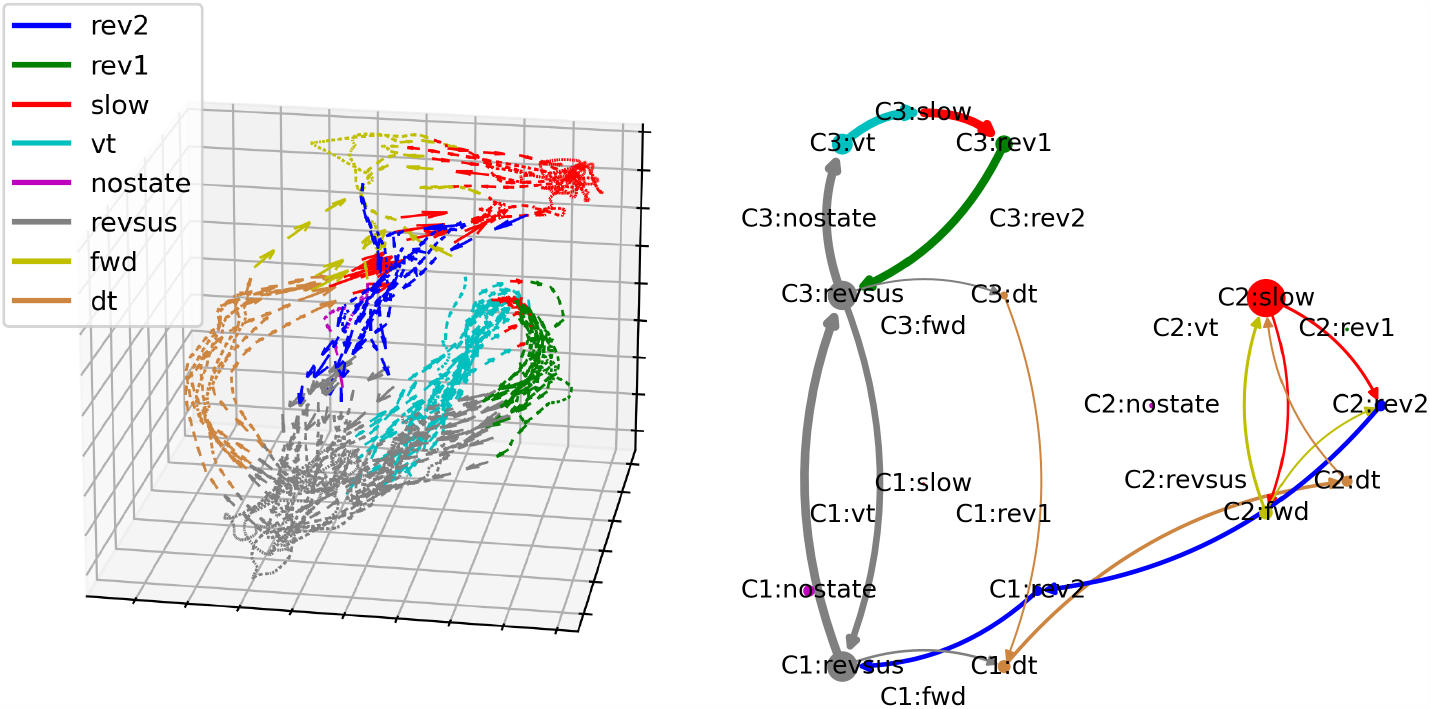
Side-by-side comparison of the neuronal manifold of the third worm as learned by BunDLe-Net and its representation as a directed graph in the NC-MCM framework.

To better understand the structure of the neuronal manifold, we now consider the branching and convergence points of the neuronal trajectories. We identify the dense cluster of neuronal states that correspond to a sustained reversal (*revsus* shown in gray at the bottom of the left plot in Figure 7) as the root node of the manifold that acts both as a branching and a convergence point. In particular, the neuronal state trajectories branch out from here into one of the two behavioral motifs via a ventral- (*vt* – turquoise) or a dorsal turn (*dt* – orange). Once a ventral turn is initiated, the neuronal trajectories exhibit almost deterministic dynamics; all trajectories execute a highly consistent loop (accompanied by a ventral turn, slowing, and reversal 1) before converging again into the root node during a sustained reversal. If a dorsal turn is initiated, on the other hand, the neuronal trajectories enter a second branching point during the subsequent slowing movement (red), from which they either return directly during a *rev2* behavior (blue) into the root node (gray) or take the longer route into the extended branch of the persistent alternation between slowing (red) and forward (yellow) behavior. Both of these branches converge again in the *rev2* behavior before returning to the sustained reversal root node. To summarize, the dynamics on the neuronal manifold are determined by a small number of branching and convergence points with highly consistent trajectories between these points.

In the cognitive-behavioral state diagram in the right column of Figure 7, the branching and convergence points of the neuronal manifold are represented by nodes with an out- and in-degree greater than one, respectively. The trajectories between the branching and convergence points are represented by nodes with an out- and in-degree of one. Based on this correspondence, it is easy to verify visually that the cognitive-behavioral state diagram constitutes a representation of the structure of the neuronal manifold as a directed graph in the sense that the nodes of the graph represent bundles of trajectories on the neuronal manifold and the edges between the nodes represent the possible paths that the neuronal trajectories can take on the manifold. This correspondence also holds for the other four worms shown in Figures 8 – 11^9^.

**Figure 8:**
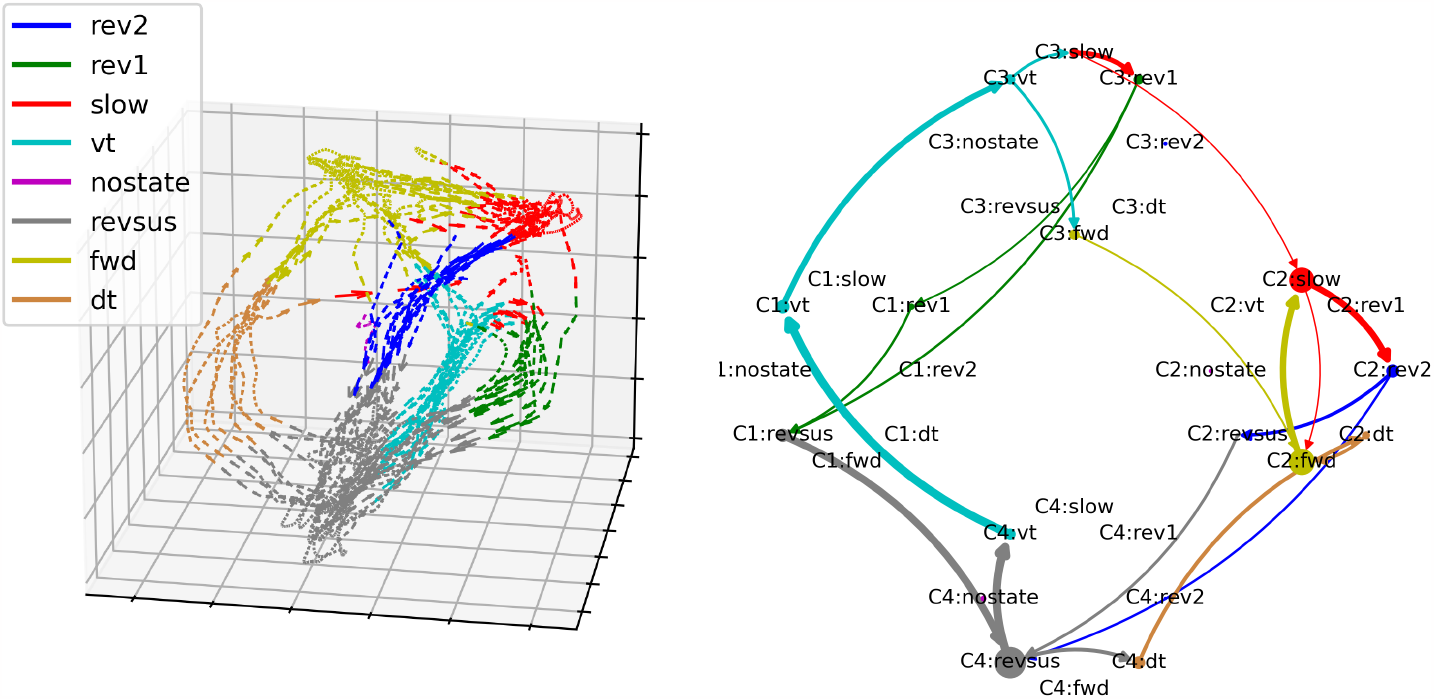
Side-by-side comparison of the neuronal manifold and the cognitive-behavioral state diagram of worm 1.

**Figure 9:**
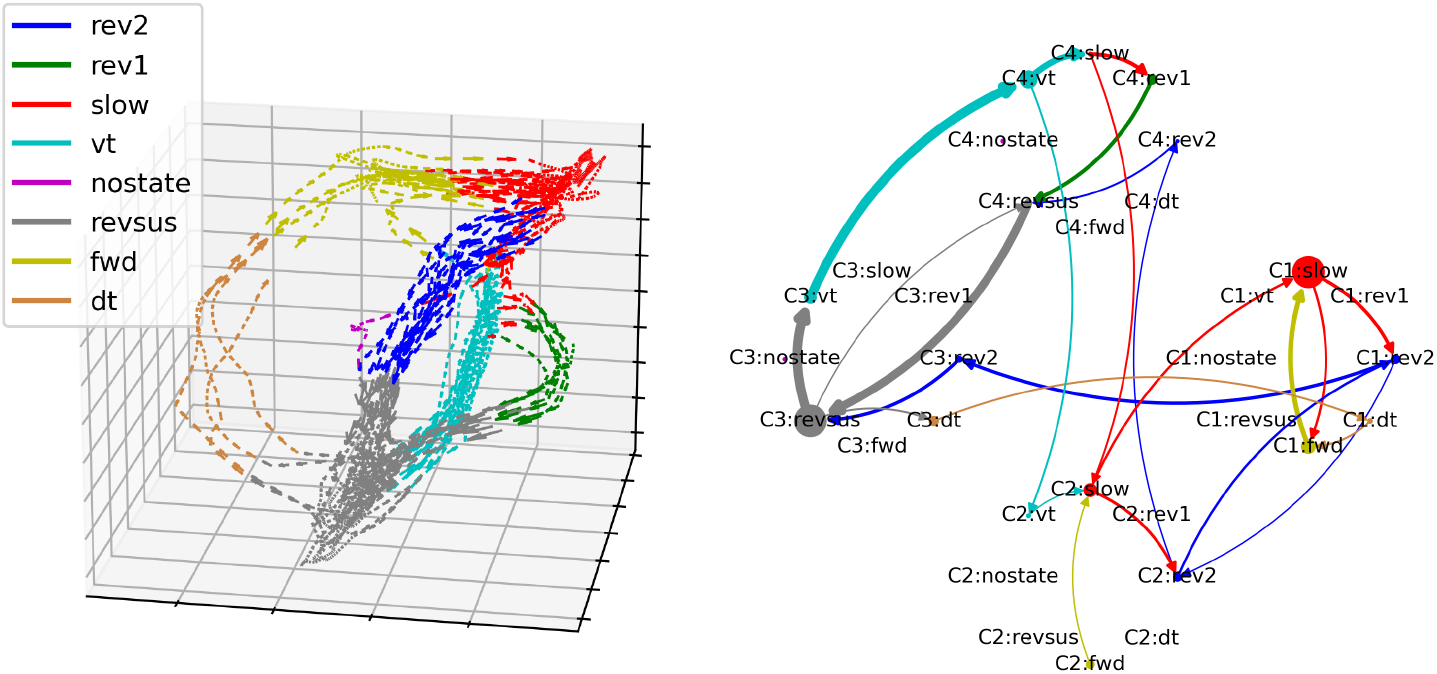
Side-by-side comparison of the neuronal manifold and the cognitive-behavioral state diagram of worm 2.

**Figure 10:**
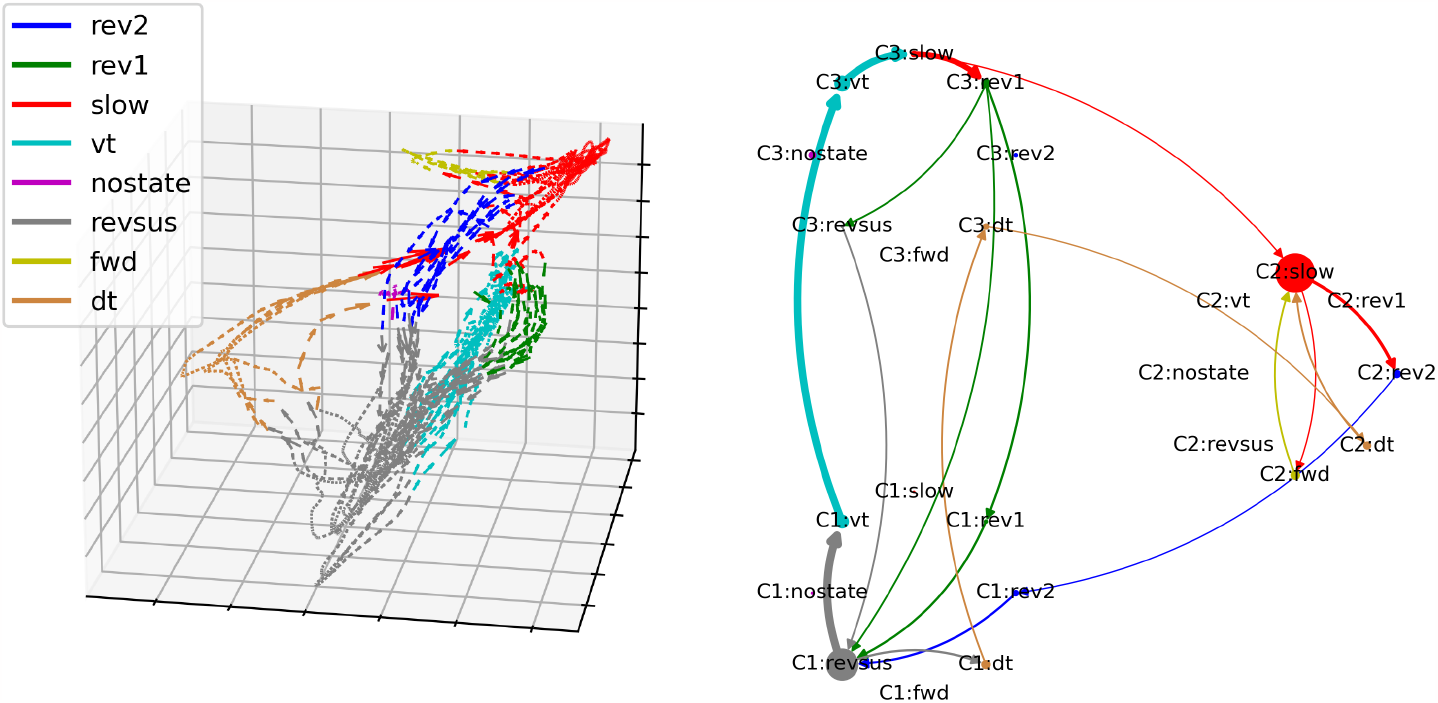
Side-by-side comparison of the neuronal manifold and the cognitive-behavioral state diagram of worm 4.

**Figure 11:**
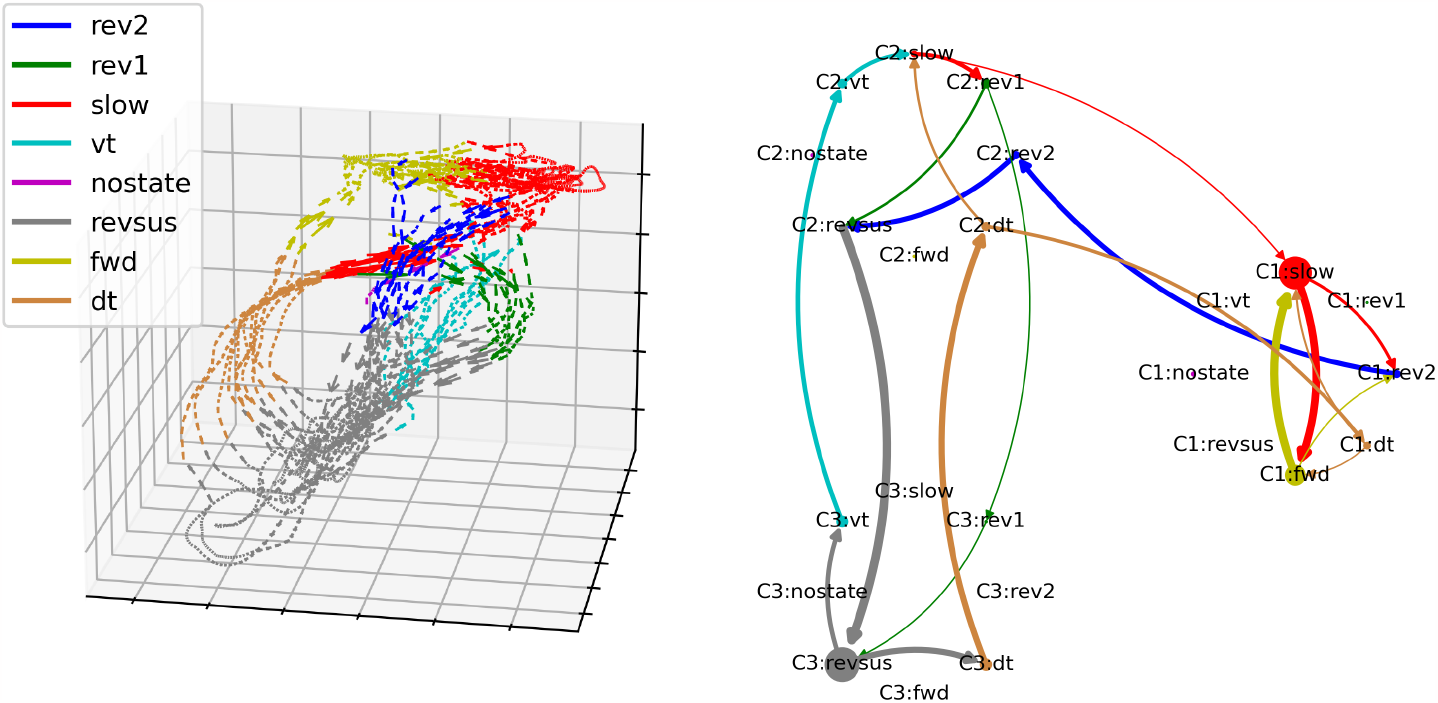
Side-by-side comparison of the neuronal manifold and the cognitive-behavioral state diagram of worm 5.

As such, the cognitive state transition models of the NC-MCM abstract away variability of the the neuronal dynamics to reveal the essential structure of the neuronal manifold in a directed graph. In this way, the NC-MCM reduces the study of the structure of neuronal manifolds to a graph theoretic problem, a mature scientific field for which a plethora of computationally efficient algorithms are available (Even, 2011).

#### 3.3.3 Towards understanding decision making in *C. elegans*

As we have seen in the previous section, the neuronal trajectories alternate between segments with almost deterministic dynamics, e.g., when executing the behavioral motif initiated by a ventral turn, and segments with high uncertainty about the future dynamics and behavior, e.g., when performing a sustained reversal movement. In the following, we show how the cognitive-behavioral state transition diagrams of the NC-MCM framework provide insights into the neuronal basis of future neuronal dynamics during segments with high uncertainty, i.e., we show how to use the NC-MCM framework to study decision-making in *C. elegans*. In particular, we study how a ventral vs. a dorsal turn is initiated during a sustained reversal and how the alternating forward and slowing behavior is terminated by a rev2-reversal.

We begin again by considering the cognitive-behavioral state transition diagram of the third worm in Figure 7. In particular, we note that the sustained reversal movement (*revsus*) is distributed across the two cognitive states C1 and C3, with frequent, cyclical transitions between the two states. Importantly, the probability that the worm transitions from a sustained reversal to a particular turning behavior differs between these cognitive states. When the worm is in cognitive state C1, it transitions from a sustained reversal into a ventral- and a dorsal turn at frequencies of 3.04% and 0.72%, respectively. In cognitive state C3, on the other hand, the frequencies of transitions from a sustained reversal into a ventral- and a dorsal turn are 0.17% and 0.84%, respectively. Ventral turns are thus roughly 18 times more frequently initiated in state C3 than in state C1 while dorsal turn initiations occur with similar frequencies in both cognitive states (they are 1.17 times more likely in C1 than in C3). We can thus interpret a cognitive state transition from C1 into C3 during a sustained reversal movement as a decision process that makes a ventral turn more likely than a dorsal turn and vice versa. We now turn to the question of how these cognitive state transitions are realized on the neuronal level.

To do so, we analyze which global perturbation of neuronal states during a particular type of movement the model predicts to induce a cognitive state that increases the probability of another type of behavior. To illustrate this idea, consider again the two states *C1:revsus* and *C3:revsus* of the third worm. By first computing the mean activation of every neuron in each of the two states, i.e., the mean of all neuronal states that map to *C1:revsus* and the mean of all neuronal states that map to *C3:revsus*, and then subtracting the two means, we obtain the neuronal perturbation that the cognitive-behavioral model predicts to induce a state transition from *C1:revsus* to *C3:revsus*; a state transition that increases the probability of transitioning into a ventral turn from 0.17% to 3.04%. To generalize from this example, we can pick a *source behavior*, e.g., a sustained reversal, and a *target behavior*, e.g., a ventral turn, determine all cognitive states in which the source behavior occurs, for each pair of cognitive states compute the neuronal perturbation that the NC-MCM model predicts to induce a transition from one to the other, weigh this neuronal perturbation by the difference in probability in the target behavior between the two cognitive states, and finally average the neuronal perturbations across all possible state transitions and worms. This results in a neuronal perturbation vector, which we subsequently denote by 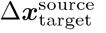, that the NC-MCM model predicts, on average across all cognitive states and worms, to induce a state transition that increases the probability of the target behavior while maintaining the current source behavior.

However, this procedure necessitates that the cognitive-behavioral state diagram of each worm represents the source behavior in multiple cognitive states. As can be checked in Figures 7 to 11, out of the cognitive-behavioral state diagrams with three cognitive states only the state diagram of the third worm represents the uncertainty during the sustained reversal movement by more than one cognitive state. This limitation can be easily remedied by considering more than three cognitive states. In the following, we consider cognitive-behavioral state diagrams with seven cognitive states, because we found seven to be the optimal number of cognitive states with respect to (the absence of evidence against) Markovianity across all worms (cf. Figure 3)^10^. Figure 12 displays the corresponding cognitive-behavioral state diagram for the third worm. Increasing the number of cognitive states from three to seven preserves the overall structure as well as the two primary behavioral motifs while splitting up the cognitive states into a more fine-grained representation of the neuronal dynamics and their relation to behavior. In particular, the pirouette motif that is initiated by a ventral turn and that was represented by cognitive state C3 in Figure 7 is represented in Figure 12 by the two cognitive states C1 and C2. The forward-run motif which is initiated by a dorsal turn and that was represented in Figure 7 by C2 is represented in Figure 12 by the two cognitive states C4 and C5. The uncertainty in the transitions between these two motifs, which was characterized in Figure 7 by the cyclical alterations between cognitive states C1 and C3 during a sustained reversal movement, is represented in the seven-states model by transitions between the cognitive states C2, C6, and C7. Moving to cognitive-behavioral state diagrams with seven cognitive states in each worm, we can thus compute the neuronal perturbation vector between a source and a target behavior across a broad range of cognitive states across all worms.

**Figure 12:**
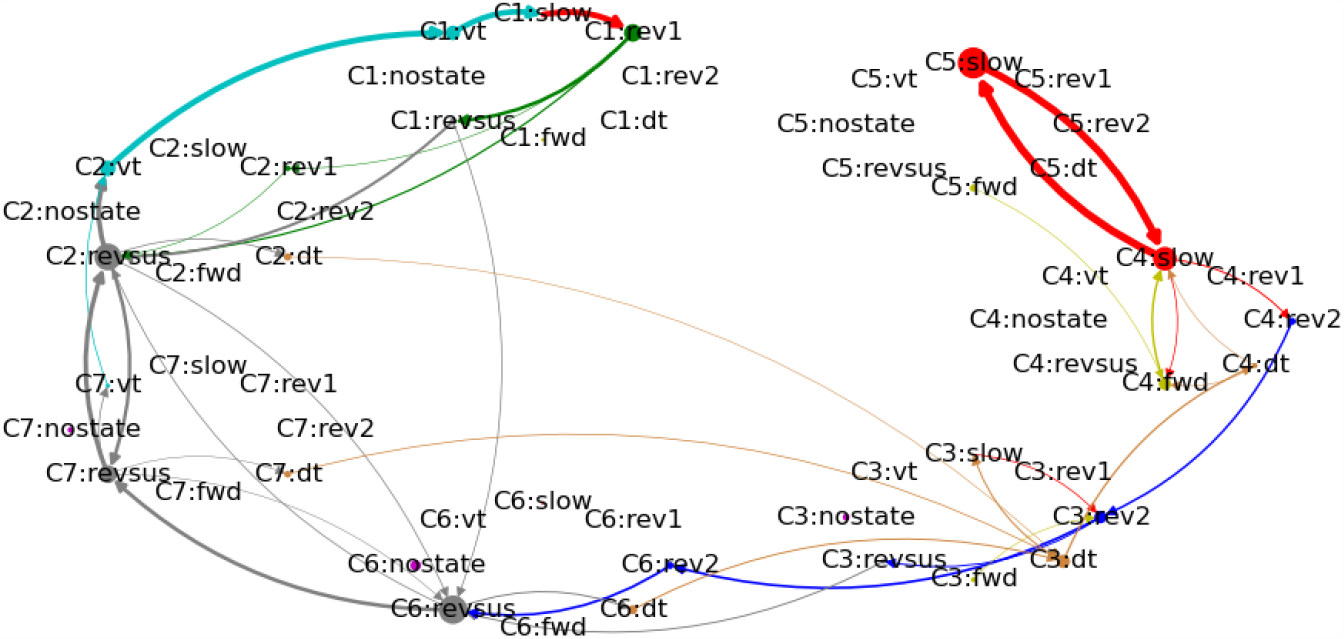
Cognitive-behavioral state transition diagram of the third worm with seven cognitive states.

Figure 13 displays the neuronal perturbation vectors 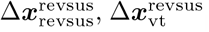, and 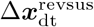 for all neurons that are shared across the recordings of all five worms, i.e., the three neuronal perturbation vectors that during a sustained reversal render the maintenance of the sustained reversal, a transition to a ventral turn, and a transition to a dorsal turn more likely. Double and single markers indicate the rejection of the null hypothesis of equal means across conditions at significance levels *α* = 0.01 and *α* = 0.05, respectively. Plus signs and asterisks indicate significance tests with and without Bonferroni correction, respectively11. Not considering those neurons used for labeling the behaviors (AVAL, AVAR, SMDDR, SMDDL, SMDVR, SMDVL, RIBR, and RIBL), the neuronal perturbation vector exhibits highly statistically significant differences across specific sets of descending interneurons. In particular, the NC-MCM model predicts that increasing the activation of AVBL and AVBL while reducing the activity of AVEL and AVER increases the probability of a turning behavior, with weaker and stronger changes linked to dorsal- and ventral turns, respectively12. Conversely, applying the opposite pattern makes it more likely that worms maintain a sustained reversal. These neuronal perturbation vectors are markedly different from those obtained for changes between cognitive states that either maintain a slowing-forward motif or interrupt this motif through a rev2-type reversal (shown in Figure 14). Here, an increase in activation of AIBL, AIBR, ASKR, RIVL, and RIVR and a decrease in activation of RMED, RMEL, and RMER are predicted to maintain a slowing behavior, with the converse pattern inducing a cognitive state that interrupts the slowing-forward motif by a rev2-type reversal.

**Figure 13:**
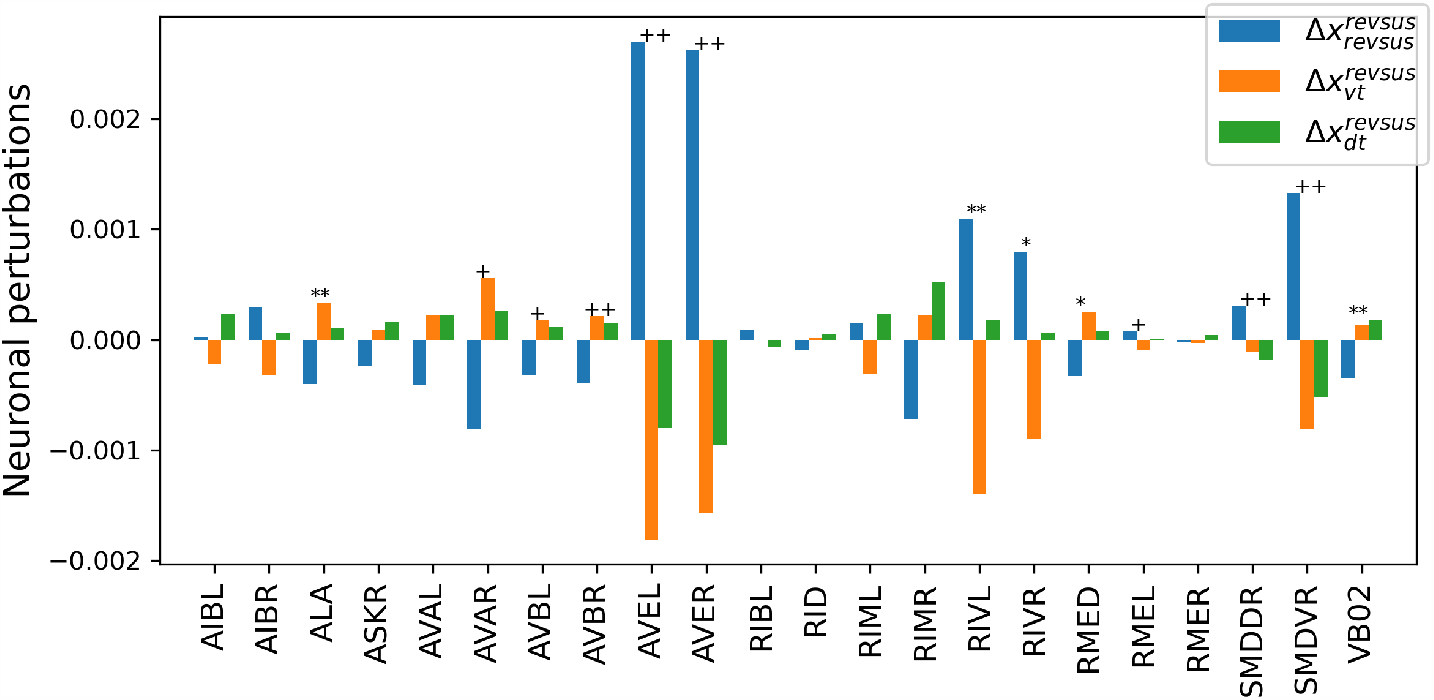
Neuronal perturbations that, when applied during a sustained reversal, the NC-MCM framework predicts to induce a cognitive state transition that increases the probability of maintaining a sustained reversal movement (blue), initiating a ventral turn (orange), or initiating a dorsal turn (green). Double and single markers indicate the rejection of the null hypothesis of equal means across conditions at significance levels *α* = 0.01 and *α* = 0.05, respectively. Plus signs and asterisks indicate significance tests with and without Bonferroni correction, respectively.

**Figure 14:**
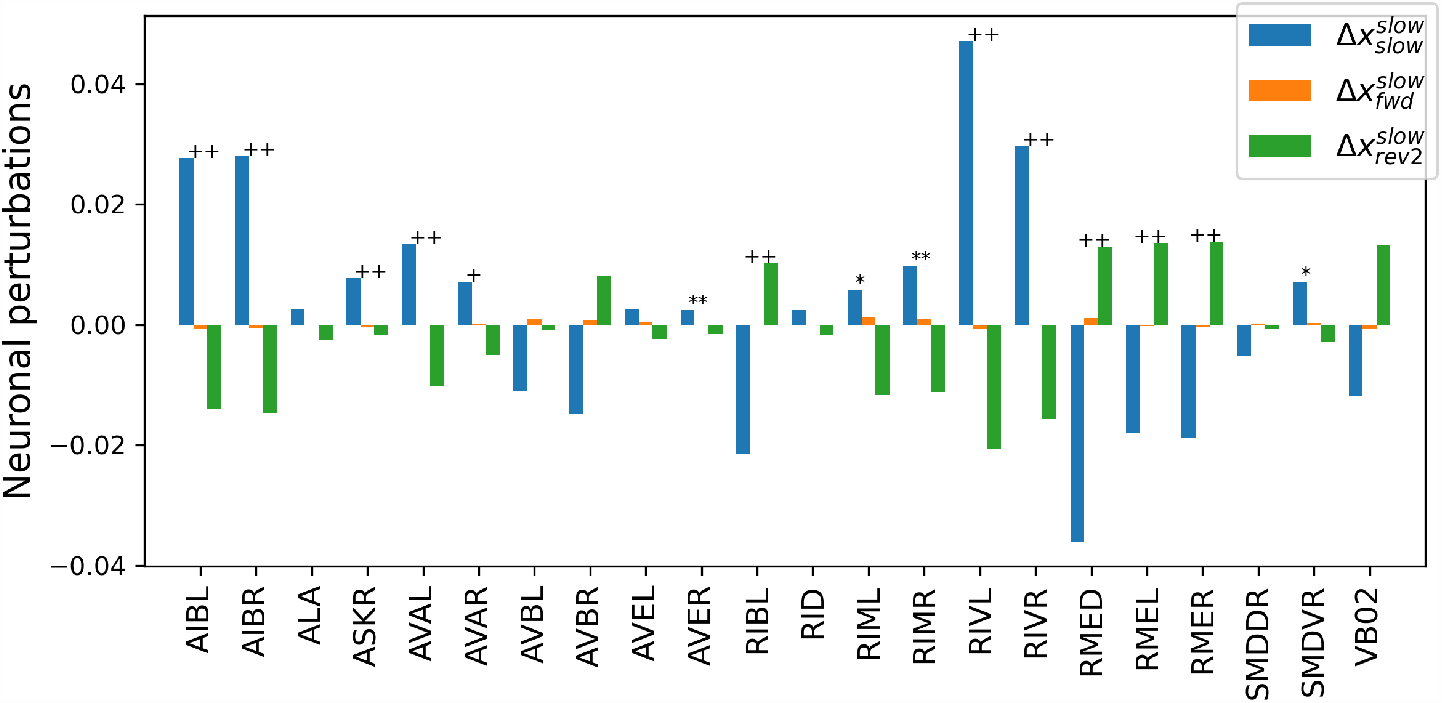
Neuronal perturbations that, when applied during a slowing movement, the NC-MCM framework predicts to induce a cognitive state transition that increases the probability of maintaining a slowing movement (blue), initiating a forward movement (orange), or initiating a rev2 reversal (green). Double and single markers indicate the rejection of the null hypothesis of equal means across conditions at significance levels *α* = 0.01 and *α* = 0.05, respectively. Plus signs and asterisks indicate significance tests with and without Bonferroni correction, respectively.

Due to the DCC property of the cognitive models discussed above, we may interpret the neuronal perturbation vectors as empirically testable predictions which perturbations of neurons via external stimulation induce cognitive states that render certain behaviors more or less likely13. However, it remains an open question how these perturbations come about naturally, i.e., how the worm decides which behavioral motif to initiate and when. A priori, there are two potential explanations. In the first explanation, the worm’s neuronal dynamics could be deterministic with the apparent randomness during sustained reversals and the slowing-forward motifs being due to some unobserved (latent) neurons. In this interpretation, observing all behaviorally relevant neurons would eliminate the randomness and lead to non-overlapping trajectory bundles in Figures 7–11. In the second interpretation, the indeterminacy may be due to inherent randomness in neuronal activation. Due to the DCC property, our results are consistent with the second but not with the first interpretation: A latent neuron that affects at least one observed neuron would induce a dependence between the current- and past states of the observed neuron, rendering the neuronal dynamics non-Markovian. Because we found no evidence against Markovianity in the cognitive state transitions, which constitute an abstraction of all behaviorally relevant information in the neuronal dynamics, our results indicate that decision-making in *C. elegans* has an intrinsically random component, as was suggested for switches between forward and backward movement (Roberts et al., 2016) 14.

### 3.4 Discussion of the experimental results in *C. elegans*

The NC-MCM framework, applied to brain wide Ca2+-imaging data of *C. elegans*, revealed cognitive states that correspond to neuronal representations of behavioral hierarchies that sit on top of a previously described motor hierarchy (Kaplan et al., 2020). A pirouette is a long time scale behavioral motif that encompasses the repetitive execution of reversal-turn command sequences, which can be triggered spontaneously or by sensory inputs (Tanimoto et al., 2017; Pierce-Shimomura et al., 1999). In our data, a pirouette manifests as a motiv of switching dynamics between neuronal activity patterns representing these actions; notably the NC-MCM framework was able to find these patterns in the absence of neuronal activities that specifically mark the initiation, duration or termination of a pirouette, i.e., which we would have interpreted as explicit upper-hierarchy neuronal representations of a pirouette. We cannot exclude that our recording technique is insensitive to such activation patterns. But alternatively, and in analogy to the pressure of a gas (see introduction), a pirouette could be seen as a useful causally consistent description of a higher-level navigational strategy of C. elegans (Tanimoto et al., 2017; Pierce-Shimomura et al., 1999) that emerges from the ‘microscopic’ dynamical switching behavior of neuronal circuits without any explicit neuronal representation. This has implication on potential control mechanisms, which, thereby could act directly at the level of synapses and neuronal excitability within these circuits, and without the need of upper-hierarchy command-circuits. In the same vein, the learned NC-MCM suggests that forward run is a dynamical motif that encompasses continuous forward crawling with intermittent slowing events. It further suggests that slowing during forward runs is neuronally disticnt from slowing during pirouettes, which we, however, lumped together into one state in our previous work (Kato et al., 2015).

When allowing for a finer granularity of many cognitive states, the NC-MCM framework can be a useful discovery tool to identify circuit elements that are potentially implicated in decision making (cf. Section 3.3.3). How *C. elegans* decides between a dorsal or ventral turn following a reversal is not known (Kato et al., 2015). The learned NC-MCM suggests circuit elements that terminate sustained reversals and influence the subsequent binary choice via expected motor neuron classes (RIV, SMDV or RMED) (Gray et al., 2005), but also unexpected neuronal cell types like the descending interneuron class AVE or the neuromodulatory neuron class ALA. Likewise, the NC-MCM makes concrete suggestion for the circuitry that controls slowing, like the AIB neurons that show Ca2+-transients during discrete slowing events in freely moving worms (Kato et al., 2015), or RIB neurons that have been implicated in the control of locomotion speed (Kato et al., 2015; Kaplan et al., 2020; Li et al., 2014). Interestingly, ASK sensory neurons, which exhibit spontaneous Ca2+-fluctuations under the experimental conditions used here (Skora et al., 2018; López-Cruz et al., 2019) are further suggested to control slowing. The NC-MCM framework, thereby, can effectively guide future interrogation strategies, such as optogenetics, to confirm the role of these classes as decison making neurons.

## 4 Discussion

The framework of NC-MCM provides a formal and mathematically rigorous framework to bridge the explanatory gap between neuronal activity and cognition. This bridging is achieved by construing cognitive states as abstractions of neuronal states that are causally consistent with respect to a set of behavioral patterns (behavioral causal consistency) and with respect to the system’s dynamics (dynamic causal consistency). As such, a NC-MCM enables us to causally reason about a system’s dynamics and behavior on the cognitive level while grounding all causal statements in the system’s neuronal states.

Dynamic causal consistency is achieved by constructing a Markovian representation of the observed dynamics. Markovianity is essential here because it guarantees a causally sufficient description of the dynamics of a physical system: Consider a simple pendulum whose complete physical state at any given point in time is given by its position and velocity. As such, setting the pendulum’s position and velocity by an external intervention fully determines its future behavior. Based on observational data only, we can check that the position and velocity provide a full characterization of the pendulum’s state by testing for Markovianity. This would reveal that the past positions and velocities do not provide additional information on the pendulum’s future states if the current position and velocity are known. Conversely, only observing the position of the pendulum does not allow us to determine its future state, e.g., because it could be swinging forward or backward15. In other words, the current position is not a Markovian representation of the pendulum’s dynamics and thus does not provide a causally sufficient representation of the physical system. Returning to the NC-MCM framework, the dynamic causal consistency property guarantees that we have found a complete, causally sufficient characterization of the neuronal dynamics that are relevant for a given behavioral context. We note that learning such a representation from empirical data is not guaranteed to succeed, e.g., because the set of observed neurons may not provide a full characterization of the organism’s physical state with respect to a given behavioral context.

In contrast to other frameworks that model cognition and its relation to neuronal dynamics, e.g., ACT-R (Anderson, 2013; Fincham et al., 2002) and atlases of cognition (Poldrack and Yarkoni, 2016; Varoquaux et al., 2018), a cognitive model in the NC-MCM framework does not incorporate prior assumptions on the structure of cognition, e.g., by postulating information-processing modules (cf. Ritter et al. (2019)) or by assuming a cognitive ontology (Varoquaux et al., 2018). Rather, a cognitive model in the NC-MCM framework is learned bottom-up from the system’s neuronal dynamics in combination with a behavioral context. In the following, we discuss in which sense such a model provides an understanding of the neuronal system it models.

On a practical level, the NC-MCM framework can be interpreted as a data compression and visualization method. By eliminating redundant information in the neuronal states with respect to the behavioral context, irrelevant information is discarded and meaningful relationships emerge in an intuitively interpretable fashion, e.g., by representing the branching and convergence points of a neuronal manifold as a directed graph. This reduction of the neuronal manifold to a directed graph opens up the rich field of graph algorithms for the study of neuronal dynamics (Even, 2011). In this view, the NC-MCM framework is a tool for cognitive neuroscientists to analyze complex neuronal data.

On a more fundamental level, we first note that the term *understanding* is not well defined and as such subjective, e.g., what qualifies as an explanation that leads to understanding by one person may be considered an insufficient explanation by another person. We adopt the viewpoint of Papineau (1998) that the deepest level of understanding is that of identities, e.g., if we learn that the morning star and the evening star are in fact the same star, i.e., Venus, it does not make sense anymore to ask why the morning star is the evening star. In analogy, the NC-MCM framework, first, describes an equivalence relation between neuronal states with respect to a set of behaviors and, second, establishes identities between the neuronal equivalence classes and cognitive states. This raises the question what a cognitive state is or what it should be. We have defined a cognitive state as a causally meaningful abstraction of a neuronal state, implying that a cognitive model captures all relevant causal relations between neuronal states and behaviors (behavioral causal consistency) and enables us to causally reason about its dynamics (dynamic causal consistency). We argue that this definition of a cognitive state is intuitively plausible because it is in line with how we use cognitive states in everyday life. Faced with the daunting task of understanding the complex neuronal systems that support our own as well as other people’s behavior, we have developed a (sometimes more and sometimes less adequate) set of concepts which we employ to reason about our own and other people’s mental processes in a given behavioral context.

We note that the granularity of the behavioral context, which is determined by the observer of a system, influences the complexity of the cognitive abstractions. For instance, only distinguishing between forward and backward movements of *C. elegans* can be expected to lead to more redundancies in the neuronal states (and hence fewer cognitive states) than labeling *C. elegans*’ movements according to angles between multiple body segments.

On a more abstract philosophical level, we remark that the NC-MCM framework offers a potential resolution to the problems of causal overdetermination and downward causation in the philosophy of mind (Woodward, 2020). By showing how a neuronal- and a cognitive level model can be constructed that are causally consistent, i.e., that allow us to interchangeably argue about a system’s dynamics and behavior on both levels, we hope that the NC-MCM framework will stimulate discussions on how to reconcile dualistic with physicalistic accounts of mental causation (Melnyk, 2003; Menzies, 2003).

Returning to more practically relevant interpretations of the NC-MCM framework, we note that (in contrast to hand-crafted mechanistic models commonly employed in computational neuroscience) the NC-MCM framework does not require mechanistic models of the neuronal-level dynamics to learn (observationally) behaviorally and dynamically causally consistent cognitive-level models. While a mechanistic model may be useful to derive the interventional distribution *P* (***B***[*t*]|do(***X***[*t*])) from which a causally behaviorally consistent model can be constructed, machine learning models in combination with a number of experimental interventions on the order of the number of cognitive states are in principle sufficient for learning *P* (***B***[*t*]|do(***X***[*t*])) (cf. Section 2.3). Being able to leverage state-of-the-art machine learning algorithms for modeling the relations between neuronal activity patterns and behavior is particularly appealing when attempting to scale up NC-MCMs from small systems such as *C. elegans*, where hand-crafted mechanistic models may be feasible, to more complex organisms consisting of potentially hundred thousands or even millions of neurons. While detailed mechanistic models may continue to provide important insights into the computations of individual circuits, we conjecture that causally consistent NC-MCMs will be more useful in the domain of cognitive neural engineering, where we would like to know how to experimentally intervene on the neuronal level of large-scale organisms to treat cognitive disorders.

To conclude the article, we discuss some extensions of NC-MCMs for future work. First, we recall that we have chosen a rather simple machine learning pipeline in Section 3 to illustrate how to learn a NC-MCM. More complex computational models of *C. elegans* are available and could be leveraged for learning NC-MCMs (Brennan and Proekt, 2019). In general, leveraging the power of state-of-the-art artificial intelligence algorithms, e.g., as being developed within the framework of causal representation learning (Schölkopf et al., 2021), is probably required to learn NC-MCMs in more complex organisms. Second, we consider the extension to multiple behavioral contexts. In Section 3, we have considered a set of behavioral states that are mutually exclusive, e.g., the worm can either crawl forward or backward but not execute both actions simultaneously. In more complex organisms, we may consider multiple behavioral contexts that are not mutually exclusive, e.g., optomotor- and swimming (bout) actions in larval zebrafish. We could then learn a NC-MCM for each behavioral context and study causal interactions between the cognitive states of each NC-MCM. Naturally, this approach could be scaled-up to an arbitrary number of behavioral contexts, potentially giving rise to a causally meaningful and neuronally grounded approach to studying cognition in complex organisms. Finally, we remark that we have focused in this work on learning cognitive states from neuronal data based on behavioral contexts. We could also consider other learning problems, e.g., settings in which sets of neuronal and cognitive states are already given and where we wish to determine whether there exists a behavioral context that gives rise to a NC-MCM. Such learning problems may be helpful to study whether already established cognitive ontologies, e.g., in psychology, are grounded in neuronal activity or, conversely, whether certain established cognitive concepts should be refined or excluded from scientific discourse. Independently of which particular learning problem we are interested in, the NC-MCM framework constitutes a rich and theoretically principled approach to bridging the explanatory gaps between neuronal activity patterns, cognitive states, and behaviors.

## Footnotes

1 We remark that this representation also allows feedback loops, e.g., as in *X*_*i*_[*t*] *→ X*_*j*_ [*t* + 1] *→ X*_*i*_[*t* + 2]

2 This test for Markovianity is implemented as the function *markovian()* in the *nilab* toolbox.

3 We remark that we could also split the data into a training- and a test set, learn and select the best clustering on the training set, and then test for Markovianity on the held-out test set. Because the number of samples available in this setting is limited, and splitting the data would further reduce the probability of finding evidence against Markovianity on the test set, we consider using all data for clustering the more conservative approach. This issue is relevant when invoking the DCC property, which we revisit in Section 3.3.3.

4 We remark again that Markovianity can not be verified. We can only fail to find evidence against Markovianity and thus accept the null hypothesis.

5 We have chosen three cognitive states because this is the lowest number of cognitive states for which we do not find evidence against Markovianity. The behavioral motifs discussed below are also apparent when considering models with more cognitive states.

6 To reduce spurious edges that result from jitters of the neuronal trajectories across the decision boundaries in the probability space, we only plot state transition if the length of a state exceeds two samples.

7 Note that the cognitive states are drawn counter clockwise with the eight behaviors of each cognitive state also arranged in circles.

8 An in-depth comparison of BundDLe-Net with other neuronal manifold learning algorithms on the present data set is available in Kumar et al. (2023).

9 For a three-dimensional visualization of the embeddings learned by BunDLe-Net, we refer to the rotating GIFs that are part of the BunDLe-Net repository at https://github.com/akshey-kumar/BunDLe-Net.

10 We remark that among all Markovian cognitive models there is no correct or incorrect number of cognitive states. We can vary the number of cognitive states and thus choose the level of granularity at which we analyze the cognitive dynamics.

11 Statistical tests were carried out with an ANOVA based on 10.000 random permutations with Bonferroni correction for multiple comparisons according to the number of neurons.

12 RIVL and RIVR may play a further role in differentiating between dorsal and ventral turns. Because these two neurons show no significant differences after Bonferroni correction for multiple comparisons, we refrain from further interpreting their effects.

13 Because the learned cognitive models are only observationally but not causally behaviorally consistent, we may not apply the same causal reasoning to the relations between cognitive state transitions and behaviors.

14 We remark that the worms have been immobilized and received no time-varying sensory input. In more ecologically realistic settings, sensory stimuli may override inherent randomness.

15 We remark that we can attempt to reconstruct the full state space by considering the pendulum’s past positions from which we infer its velocity.

